# AutoMorFi: Automated Whole-image Morphometry in Fiji/ImageJ for Diverse Analyses and Discoveries

**DOI:** 10.1101/2024.07.26.605357

**Authors:** Ouzéna Bouadi, Chenkai Yao, Jason Zeng, Danielle Beason, Nyomi Inda, Zoe Malone, Jonathan Yoshihara, Amritha Vinayak Manjally, Clifton Johnson, Jonathan Cherry, Chin-Yi Chen, Tzu-Chieh Huang, Bogdana Popovic, Maria Henley, Guangmei Liu, Hannah Aichelman, Sarah W. Davies, Yuan Tian, Hengye Man, Thomas Gilmore, Elif Ozsen, Kristen Harder, Peter Walentek, Elizabeth K. Kharitonova, Ella Zeldich, David Pitt, Tuan Leng Tay

## Abstract

Running on the highly popular and accessible ImageJ/Fiji platform for biological image analysis, we have established AutoMorFi as a streamlined interface for automated whole-image morphometric analysis that generates at least 47 measurements per cell or object in under 1 minute. We performed multiple validated cluster and principal component analyses on nonredundant morphometric parameters derived from AutoMorFi for various cell types, objects, and organisms. We used images of rodent macrophages, human postmortem brain tissues from multiple sclerosis (MS) and Alzheimer’s disease (AD) patients, iPSC/animal models for Down’s syndrome and autism spectrum disorder (ASD), and organisms such as sea anemone and corals. AutoMorFi’s adaptability extends across diverse imaging modalities including brightfield, confocal, or widefield fluorescence microscopy as well as underwater photography. Due to its unlimited and unbiased sampling across any image and high potential for modification and customization, using AutoMorFi has led to the discovery of new distinguishing features in previously studied cell types and organisms as well as the development of rapid diagnostic approaches. AutoMorFi represents a transformative tool that will accelerate morphometric analysis and offer broad relevance in biological studies.

## MAIN TEXT

Morphometry is a bedrock for understanding life sciences. As a key indicator of functional, behavioral, and physiological state, the study of cellular and organismic morphology has been achieved through numerous methods. Common approaches range from meticulously hand-drawn neurons and camera lucida traces of glia^1–3^ to computational and machine-learning-based image analysis tools that classify cells and animal species by their diverse characteristics^4–6^. Current semi-automatic processes for common applications such as cell detection, quantification, and morphometry are, however, mostly out of reach for many researchers who are unfamiliar with coding languages that may pose a steep learning curve^7^. The day-to-day reality of image data analysis involves manual adjustments of brightness, contrast, and thresholds for each image as well as painstaking tracing and validation of individually segmented cells or objects. Accurate execution of these routine tasks requires varying in-depth subject knowledge and depends on the complexity and proximity of the objects of interest. Ideally, morphometric analysis tools should be user-accessible, generate accurate results that can be easily quality controlled, offer options for unbiased and batch sampling, and highly versatile and customizable to address multifaceted scientific questions.

For nearly 3 decades, ImageJ or Fiji^8,9^ has been the go-to open-source platform for researchers globally to process and analyze images as well as to reconstruct cells and objects acquired by microscopy to obtain quantitative morphometric data. Taking advantage of this familiar resource, we aimed to standardize and automate the acquisition of multiple morphological parameters of complex and branched cells such as the highly ramified brain-resident microglia during homeostasis, by looping a series of Fiji commands using a macro^10^. We invited 30 users ranging from high school interns to principal investigators to test our first prototype, “AutoGliji”, on a standard fluorescence image of IBA-1-immunolabeled mouse microglial cells (**Fig. 1A**). Testers were asked to perform manual measurements of the same 3 cells using Analyze Particles and the Simple Neurite Tracer ImageJ plugin^11^ for comparison (**Fig. 1A-C**). The time needed to analyze all cells in the image by AutoGliji (5.5 ± 0.75 minutes) was drastically reduced when compared to manual measurement of 3 cells (105 ± 19 minutes) (**Fig. 1D**). Furthermore, manual measurements of over half of the parameters were highly inconsistent across testers due to different levels of familiarity with microglial morphology, in stark contrast to the reproducible results from AutoGliji which were generated by the same users running either Windows or MacOS (**Fig. 1E-N**).

**Figure 1:**
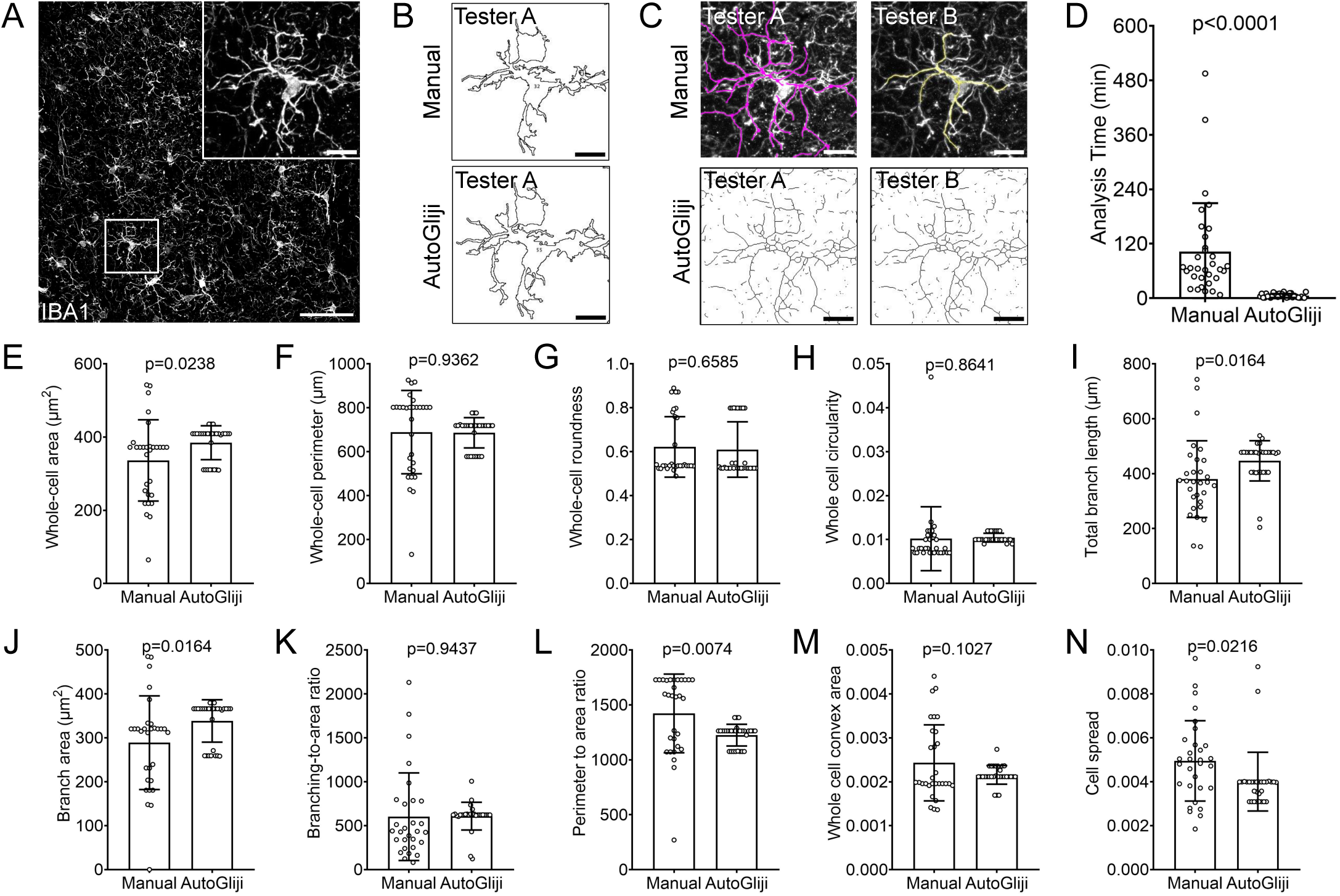
Acquisition and validation of whole-image cell morphometric data using the AutoMorFi prototype, AutoGliji. **(A)** Widefield fluorescence image of IBA-1-immunolabeled mouse cortical microglia (white) used for manual and automated morphometric analysis of 3 selected cells by 30 testers. Inset shows cell #3 corresponding to data in **B-N**. **(B)** Representative manual and automated cell thresholding and detection by 1 tester. **(C)** Representative manual trace using Simple Neurite Tracer and automated cell traces using AutoGliji by 2 testers. **(D)** Total time taken per tester for manual and automated data acquisition for cells #1–3. **(E-N)** Examples of manual and automated morphometric measurements of cell #3 for **(E)** whole-cell area, **(F)** whole-cell perimeter, **(G)** whole-cell roundness, **(H)** whole-cell circularity, **(I)** total branch length, **(J)** branch area, **(K)** branching to area ratio, **(L)** perimeter to area ratio, **(M)** whole cell convex area, and **(N)** cell spread. **(D-N)** Data are represented as mean ± s.e.m. Paired t-test. *N* = 30. Scale bars, 100 µm (overview in **A**); 10 µm (inset in **A**, **B**, and **C**).

AutoMorFi is an optimized version of AutoGliji that was configured to generate 33 measurements and 14 calculated parameters per cell or object with macro runtime reduced to under 1 minute per image (**Table 1, Supplementary Notes 1–6**, **Supplementary Video 1**). Sampling limit within any 2-dimensional 8-bit TIFF image that could be opened in Fiji was eliminated. To validate our hypothesis that 47 measures provide sufficient dimensionality to distinguish and classify cells based on morphology using standard clustering algorithms for high dimensional data, we analyzed 393 microglia, 438 cardiac macrophages, 201 alveolar macrophages, 307 liver Kupffer cells, 332 kidney macrophages, and 274 splenic macrophages from the same wildtype adult mice that were immunolabeled with IBA-1^12–14^ (**Fig. 2A-B**). As expected, 47 morphometric features classified microglia and cardiac macrophages as most morphologically distinct and revealed that macrophages of the kidney, liver, lung, and spleen share an intermediate phenotype (**Fig. 2C-F**). Furthermore, UMAP visualization of the data has allowed us to determine the major morphological characteristics that distinguish these macrophages of common lineage (**Fig. 2G-J)**.

**Figure 2:**
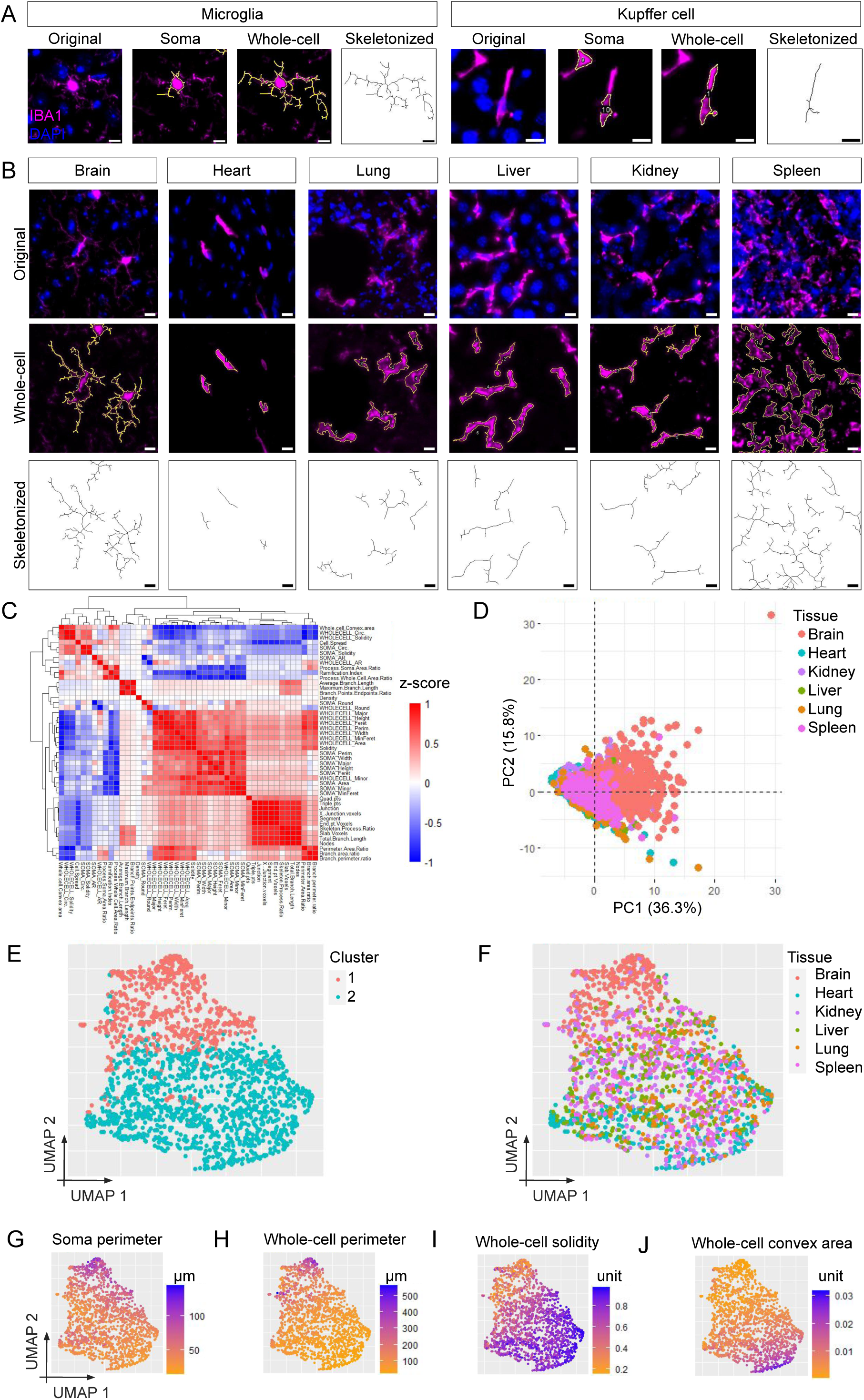
Morphometric analysis of tissue-resident macrophages. **(A)** Representative widefield fluorescence images of IBA-1^+^ brain-resident microglia and liver Kupffer cells and corresponding elements generated by AutoMorFi including soma outline, whole-cell outline, and skeletonized structure of the whole-cell outline. **(B)** Representative images of homeostatic mouse IBA-1^+^ tissue-resident macrophages and their corresponding whole-cell outlines and skeletonized traces. **(C)** Correlation matrix of the 47 parameters from AutoMorFi across 1,945 tissue-resident macrophages. **(D)** Principal component analysis (PCA) of 1,945 macrophages based on 47 parameters from AutoMorFi. **(E)** Uniform manifold approximation and projection (UMAP) of 1,945 macrophages in 2 clusters based on 24 morphometric parameters after dimensionality reduction and unsupervised kmeans clustering. **(F)** UMAP of macrophages according to tissue origin. **(G-J)** UMAP of selected parameters, including **(G)** soma perimeter, **(H)** whole-cell perimeter, **(I)** whole-cell solidity, and **(J)** whole-cell convex area. Scale bars, 10 µm.

**Table 1.** List of 47 morphometric parameters.

| Category | Parameter | Definition |
| --- | --- | --- |
| "Soma" | 1. Soma area | Area of the object (square micrometers) |
|  | 2. Soma perimeter | Length of the outside boundary of the object (micrometers) |
|  | 3. Soma width | Width of the smallest rectangle enclosing the object (micrometers) |
|  | 4. Soma height | Height of the smallest rectangle enclosing the object (micrometers) |
|  | 5. Soma major | Primary axis of the best fitting ellipse in the object (micrometers) |
|  | 6. Soma minor | Secondary axis of the best fitting ellipse in the object (micrometers) |
| | 7. Soma circularity | Value of 1.0 indicates a perfect circle, $4\pi \cdot \text{area} / \text{perimeter}^2$ |
|  | 8. Soma feret | Maximum caliper, longest distance between any two points in the object |
|  | 9. Soma minimum feret | Minimum caliper diameter |
|  | 10. Soma aspect ratio | Major axis/minor axis |
| | 11. Soma roundness | Inverse of the aspect ratio, $4 \cdot \text{area} / (\pi \cdot \text{major axis}^2)$ |
|  | 12. Soma solidity | Area / convex area |
| Whole-cell | 13. Whole cell area | Area of the object (square micrometers) |
|  | 14. Whole cell perimeter | Length of the outside boundary of the object (micrometers) |
|  | 15. Whole cell width | Width of the smallest rectangle enclosing the object (micrometers) |
|  | 16. Whole cell height | Height of the smallest rectangle enclosing the object (micrometers) |
|  | 17. Whole cell major (X) | Primary axis of the best fitting ellipse in the object (micrometers) |
|  | 18. Whole cell minor (Y) | Secondary axis of the best fitting ellipse in the object (micrometers) |
| | 19. Whole cell circularity | Value of 1.0 indicates a perfect circle, $4\pi \cdot \text{area} / \text{perimeter}^2$ |
|  | 20. Whole cell feret | Maximum caliper, longest distance between any two points in the object |
|  | 21. Whole cell minimum feret | Minimum caliper diameter |
|  | 22. Whole cell aspect ratio | Major axis/minor axis |
| | 23. Whole cell roundness | Inverse of the aspect ratio, $4 \cdot \text{area} / (\pi \cdot \text{major axis}^2)$ |
|  | 24. Whole cell solidity | Area / convex area |
| "Skeletonized" | 25. Total number of segments | Total number of slab segments |
|  | 26. Number of junctions | Number of actual segment junctions |
|  | 27. Number of end point voxels | Voxel with less than 2 neighbors |
|  | 28. Number of junction voxels | Voxel with more than 2 neighbors |
|  | 29. Number of slab voxels | Voxel with exactly 2 neighbors |
|  | 30. Average branch length | Average length of branches (micrometers) |
|  | 31. Number of triple points | Number of junctions with exactly 3 branches |
|  | 32. Number of quadruple points | Number of junctions with exactly 4 branches |
|  | 33. Maximum branch length | Maximum length of branches (micrometers) |
| Additional parameters | 34. Total branch length | Maximum branch length / Number of junction voxels |
|  | 35. Cell spread | Whole cell aspect ratio / Number of end point voxels |
|  | 36. Branch area ratio | Whole cell area / Soma area |
|  | 37. Branch perimeter ratio | Whole cell perimeter / Soma perimeter |
|  | 38. Whole cell convex area | Whole cell solidity / Whole cell area |
|  | 39. Nodes | Number of end point voxels + Junction voxels + Slab voxels |
| | 40. Ramification index | $\text{Whole cell perimeter} / \text{Whole cell area} / 2 \times [\sqrt{(\pi \cdot \text{Whole cell area} \times XY)}]$ |
| | 41. Perimeter to area ratio | $\text{Whole cell perimeter}^2 / \text{Whole cell area}$ |
| | 42. Density | $\text{Whole cell area} / (\text{Whole cell major} \times \text{Whole cell minor})$ |
|  | 43. Process soma area ratio | Branch area / Soma area |
|  | 44. Process whole cell area ratio | Branch area / Whole cell area |
| | 45. Skeleton to process ratio | $\text{Total branch length}^2 / \text{Branch area ratio}$ |
|  | 46. Branch points endpoints ratio | Number of slab voxels / Number of end point voxels |
|  | 47. Whole cell solidity | Whole cell area / Whole cell convex area |

To assess the potential of AutoMorFi for diagnostic and future large-scale applications, we tested its ability to cluster microglia identified by a single IBA-1 marker in well-documented lesions of MS from patients and controls (**Table 2**, **Fig. 3**). Microglia play different roles in all stages of demyelination, oligodendrocyte death, and neuroaxonal injury in MS^15–18^. While microglia exhibit degenerative and pro-inflammatory features as MS progresses, they also participate in neuroprotection and repair. Thus, exploring microglial morphological diversity in different lesions may unveil new strategies against microglia-driven neurodegeneration in MS. Cluster analysis revealed a clear transition of microglial morphology from control white matter (WM) to the rim of chronic active lesions (**Fig. 3C-F**) and 11 key morphological features that promoted optimal classification of the pathological stages (**Fig. 3G-J**). AutoMorFi provides a cost-effective and time-efficient alternative as a preliminary diagnostic tool with shorter turnaround time in contrast to requiring multiple immunohistochemical assays on several disease markers. Furthermore, evaluation of AutoMorFi against well-established open-source segmentation tools, including CellProfiler, ilastik, Cellpose, and QuPath suggests that only AutoMorFi offers high confidence in capturing intricate characteristics of complex and densely distributed cells with the operational efficiency necessary for handling large datasets.

**Figure 3:**
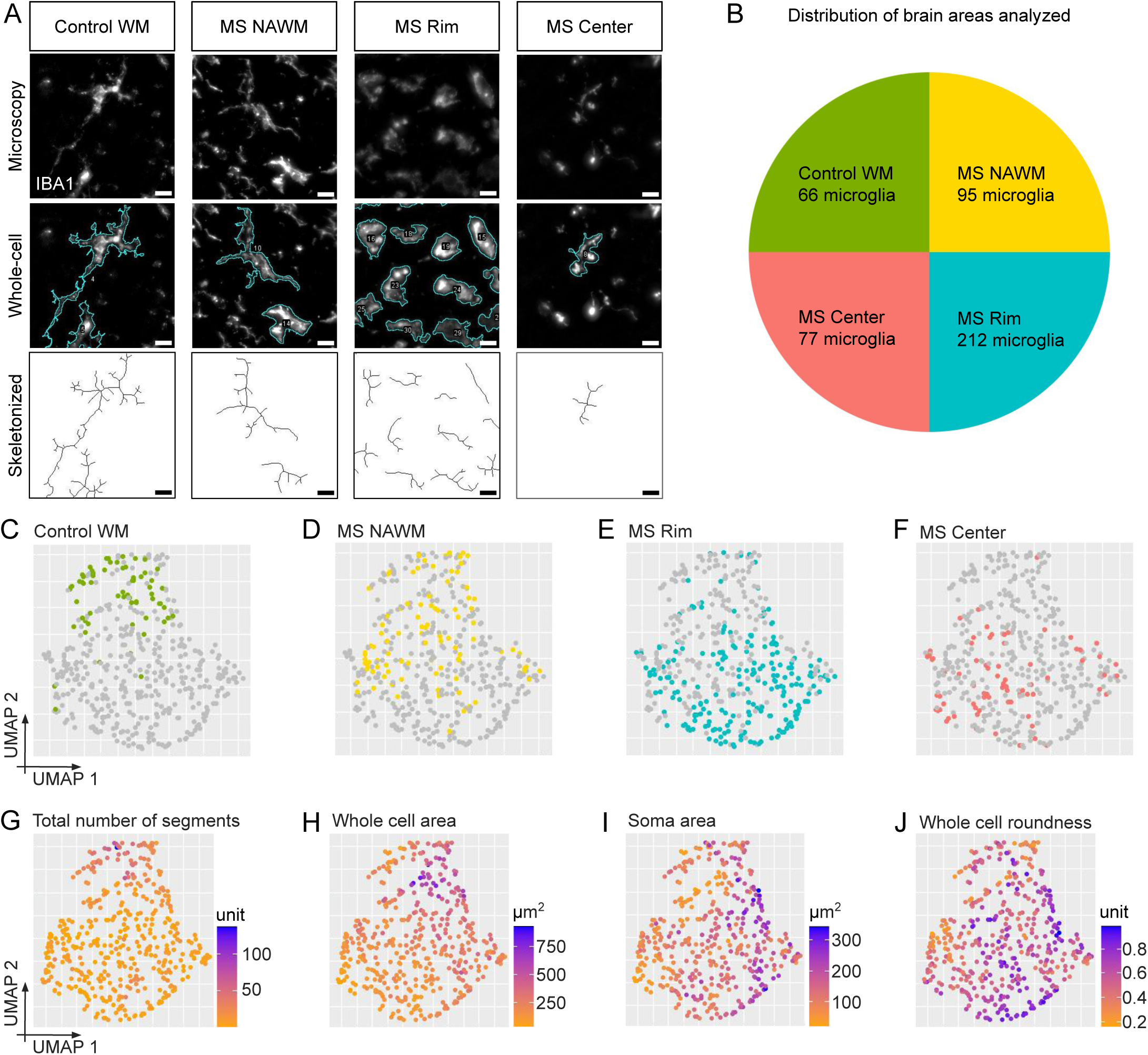
Microglial morphometric analysis in postmortem human MS and control brain. **(A)** Representative widefield fluorescence images of IBA-1^+^ microglia (white) and their corresponding whole-cell outline and skeletonized trace. **(B)** Microglial cells analyzed from the 4 equally sampled brain areas: control white matter (WM), MS normal-appearing WM (NAWM), MS rim, and MS center. **(C-J)** UMAP of 450 microglia based on 27 morphometric parameters after dimensionality reduction from **(C)** control WM, **(D)** MS NAWM, **(E)** MS rim, and **(F)** MS center and of selected parameters including **(G)** total number of segments, **(H)** whole cell area, **(I)** soma area, and **(J)** whole cell roundness. Scale bars, 10 µm.

**Table 2.** Morphometric analysis of IBA-1-immunostained microglia in postmortem human brain tissue.

| Group | Patient ID | Area analyzed (µm<sup>2</sup>) | Mean NND ± s.e.m. (µm) |
| --- | --- | --- | --- |
| Control WM | Control_56BT1, Control_56B9F, Control_31B1 | 320,000 | 41.69 ± 2.48 |
| MS NAWM | MS_590939, MS_560L2 | 320,000 | 36.41 ± 1.93 |
| MS Center | MS_590939, MS_560L2, MS_590937 | 320,000 | 38.62 ± 4.37 |
| MS Rim | MS_590939, MS_560L2, MS_590937 | 320,000 | 27.42 ± 1.87 |

WM, white matter. MS, multiple sclerosis. NAWM, normal appearing WM.

As further proof-of-principle that AutoMorFi can be easily applied for morphometry-based discoveries, it was used to analyze diverse image types from widefield or confocal fluorescence and brightfield microscopy to underwater photography (**Fig. 4**). These images represent diverse applications such as the comparison of glial cells across model systems such as euploid human induced pluripotent stem cells-derived microglia from Down’s syndrome patients, aged human microglia, *Drosophila* glia, and rodent microglia (**Fig. 4A**), characterization of *in vitro* neuronal cultures (**Fig. 4B**), single cells (**Fig. 4B-C**), or whole organisms (**Fig. 4D**), genetic studies (**Fig. 4B-C**), and the impact of environmental stress on immune signaling and coral reef ecosystems (**Fig. 4D-E**). AutoMorFi measurements unveiled additional morphological changes compared to freehand tracing^19^ of Ube3a 2x Tg neurons which model Ube3a overdosage and hyperactivity in ASD^20–24^ and confirmed the hypothesis of reduced dendritic arborization^25–27^ (**Fig. 4B**). Motile cilia are membrane-covered cell protrusions found in ciliated cells involved in facilitating movement or transport of substances. Impaired ciliated cells can lead to chronic airway diseases, infertility, or hydrocephalus. Ciliogenesis requires precise molecular regulation at the centriole to switch from cell division to cilia formation employing molecules such as the centriolar protein Cp110^28^. Fast readouts from multiplexing AutoMorFi and fluorescence intensity measurements from Centrin4 and acetylated α-tubulin markers of multi-ciliated cells (MCCs) validated previous findings of more efficient rescue of *Cp110* knockdown by *Cp110-τ<3’UTR* DNA than *Cp110* DNA (**Fig. 4C**). This approach revealed the heterogeneity of MCC phenotypes in the study and identified 30% of the parameters that were significantly altered. Starvation of sea anemones was reported to reduce NF-κB, leaving the animals vulnerable to infection^29^. NF-κB proteins are important for many cellular processes including immunity, inflammation, development, cellular growth, and apoptosis. They are differentially regulated in many diseases such as cancer, arthritis, chronic inflammation, asthma, neurodegenerative disease, and heart disease. By reanalyzing the images of whole-mount anemone polyps^29^ using AutoMorFi as well as taking intensity and distribution (area fraction) measurements of NF-κB, we newly discovered that starvation does not alter the gross morphology of the anemones and validated the earlier findings of NF-κB reduction (**Fig. 4D**). Coral reefs are critically threatened by climate change. Recent advancements in sequencing for non-model systems have revealed previously unrecognized ‘cryptic coral lineages’ (*i.e.,* genetically distinct but morphologically similar taxa) that have important implications for predicting reef ecosystem futures and effectively restoring reefs^30^. Rapid and cost-effective methods to identify these cryptic lineages, preferably alternative methods to DNA sequencing, are needed to effectively characterize coral diversity present on reef ecosystems and ultimately improve lineage assignment. AutoMorFi identified diagnostic skeletal phenotypes and replicated the same pattern of cryptic lineage differentiation in much less time compared to previous analysis requiring ∼25 minutes per image to measure all relevant traits (**Fig. 4E**).

**Figure 4:**
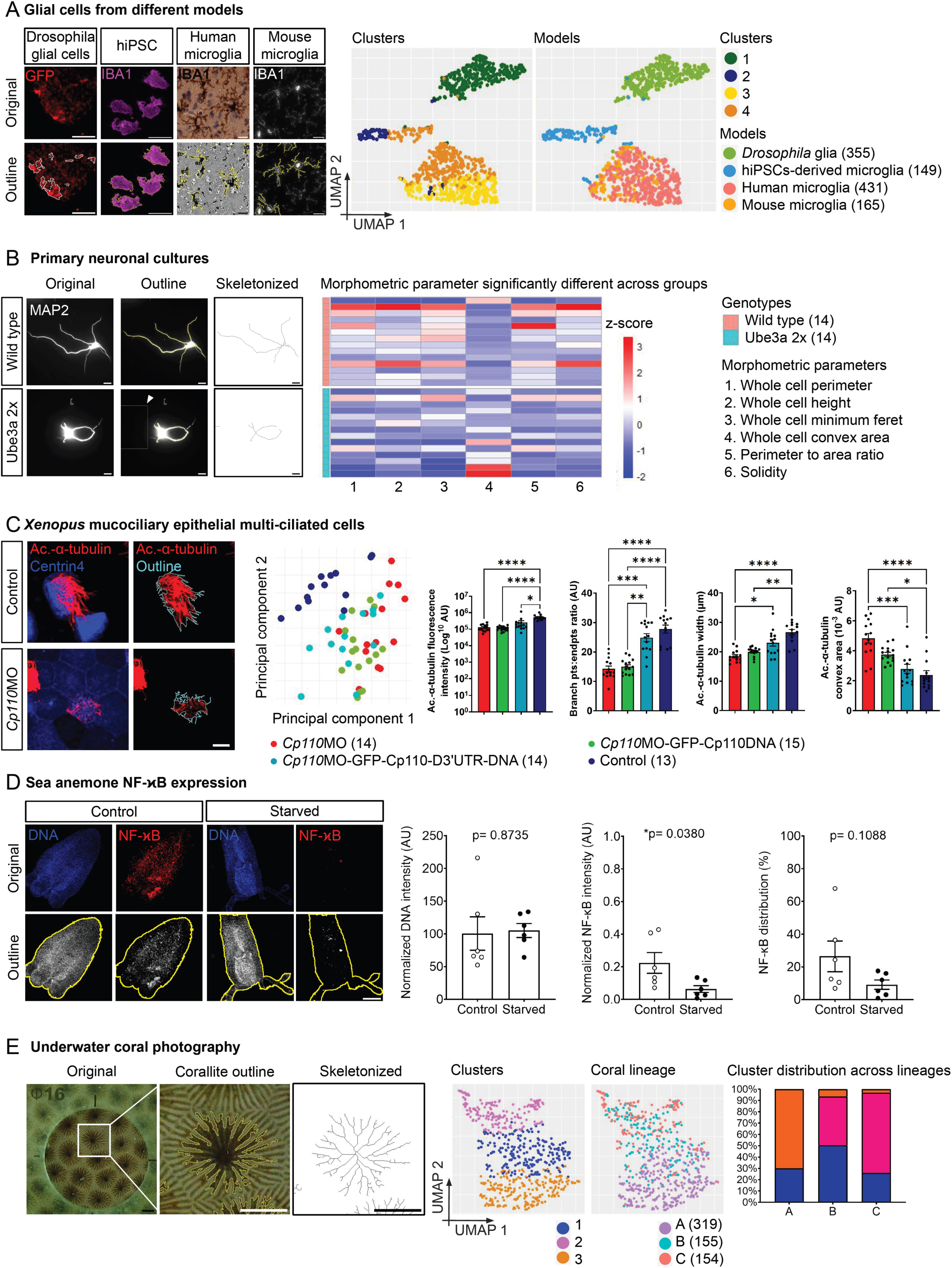
Diverse applications of AutoMorFi. (A) Comparative morphometric analysis of glial cells from different models. Representative widefield fluorescence and brightfield images of glial cells from experimental models (*Drosophila* astrocyte-like glia, hiPSCs-derived microglia, human microglia, and mouse microglia) and their corresponding whole-cell outlines. UMAPs of 1,100 glial cells clustered based on 47 morphometric parameters or model system. **(B) Single object morphometric analysis.** Representative widefield fluorescence images of MAP2-immunolabeled primary neurons (white) from Ube3a 2x transgenic and wildtype mice and their corresponding whole-cell and skeletonized traces with an example of an added pseudo-object (white arrowhead). Six of 33 morphometric parameters were significantly different between genotypes in unpaired t- tests: whole cell perimeter (p=0.0138), whole cell height (p=0.0270), whole cell minimum feret (p=0.0116), whole cell convex area (p=0.0177), perimeter to area ratio, and solidity (p=0.0351). *N* = 14 per group. **(C) Acetylated- α-tubulin marker-specific morphometric and fluorescence intensity measurements.** Representative confocal images of targeted *Xenopus* MCCs indicated by centrin4-cfp (blue) with varying ciliation indicated by acetylated-α-tubulin (red) and whole-cell outline of the marker (cyan). PCA based on 43 measurements of targeted MCCs averaged per individual embryo (N) in *Cp110* knockdown (red), rescue (green, turquoise), and control (blue) groups. Representative histograms (mean ± s.e.m.) of features with significant changes in ciliation analyzed by Kruskal-Wallis and Dunn’s posthoc tests. **(D) Whole organism morphometric and NF-κB marker analyses.** Representative widefield fluorescence images of control and starved sea anemone polyps. DNA (blue), NF-κB (red), and AutoMorFi generated whole-object outline (yellow). Selected measurements (mean ± s.e.m.) are shown. Unpaired t-tests. *N* = 6 per group. **(E) Whole-image 2D morphometric analysis of photographed objects.** Representative image of corallites photographed under water and a corresponding whole-object outline and skeletonized trace. UMAPs of 628 corallites based on 21 morphometric parameters after dimensionality reduction distributed into clusters and lineages A–C. Histogram illustrates the proportion of the 3 clusters across lineages. Scale bars, 10 µm **(A-C)**; 100 µm **(D)**; 2 mm **(E)**.

In summary, we demonstrate the easy usage and versatility of AutoMorFi for rapid and reproducible acquisition of large morphometric datasets across biological scales and research objectives that have expanded the depth of information resulting in new discoveries. Using AutoMorFi, statistical power is easily increased due to unlimited and unbiased sampling. To promote broad usage and customization of the tool for novel applications, we provide a troubleshooting guide (**Supplementary Note 3**) of common pitfalls identified from thorough executions of AutoMorFi by multiple research teams. AutoMorFi could, for example, be adapted to quantify and classify condensates of cellular proteins undergoing liquid-liquid phase separation to introduce more quantitative descriptors of proteostasis in disease and aging to the field^31^. When applied to multi-channel images (**Fig. 4C-D**) and combined with additional approaches such as genetics and genomics as well as other classification algorithms (e.g., random forest, bootstrapping^5^), morphometrics will serve as potent indicators and predictors of disease states and environmental changes.

## ACKNOWLEDGEMENTS

The authors thank AutoGliji testers E. Batchelor, S. Brayton, V. Chiplunkar, E. Choi, S. Guerra, K. He, A. Hoo, M. Iordanov, S. Jo, T. Keefauver, J. Koo, E. Kraft, H. LeBlanc, A. Liu, E. Maloney, C. Reddy, P. Ruiz, K. Sampat, and M. Zreik; I. Amin for assistance with microscopy; G. Lin (R scripts) and A. Labadorf for data analysis and visualization advice. *In memoriam* M. Spelleken and Scarlet. TLT was supported by a NARSAD Young Investigator Grant from the Brain & Behavior Research Foundation, Spivack Neuroscience Pilot Award, and Patricia McLellan Leavitt Research Fund Award.

## AUTHOR CONTRIBUTIONS

ZM, DB, CY, and OB established the macro. CY wrote the coordinate-matching code. OB, JZ, CY, NI, JY, AVM, CJ III, JC, C-YC, T-CH, BP, MH, GL, EKK, EZ, HA, SD, PW, YT, HM, TG, CHC, B-BK, DP, and TLT generated data, provided/validated images, and performed analyses. EO and KH ran pilot analysis. OB and TLT wrote the manuscript. TLT conceived and supervised the study.

## MATERIAL AND METHODS

### Sample preparation

#### Mouse

Female C57Bl/6J (wild-type) mice were purchased from Charles River Laboratories and raised in a specific-pathogen-free facility under 10h/14h dark/light cycle until 52 weeks old. Mice were given food and water *ad libitum*. Animal experiments are approved by the Institutional Animal Care and Use Committee (IPROTO202200000054). Mice were deeply anesthetized by intraperitoneal administration of 100 mg ketamine and 5 mg xylazine per kg body weight and transcardially perfused with phosphate-buffered saline (PBS). Brains, hearts, kidneys, livers, lungs, and spleens were post-fixed overnight at 4°C in 4% paraformaldehyde (PFA) in PBS, rinsed in PBS, and cryoprotected in 30% sucrose/PBS at 4°C until fully submerged. Samples were embedded in Tissue-Tek O.C.T. compound (Sakura Finetek, USA) for frozen sectioning on a cryostat (Cryostar NX50; Epredia Cryostat, USA).

#### Human

Human postmortem brain tissues from 2 MS and 2 control patients were obtained according to an institutional review board-approved protocol (Yale HIC 1211011031) (**Table 2**).

Brain hemispheres were fixed with formalin for a minimum of 2 weeks. Brain slabs with lesions were embedded with paraffin. A chronic active MS lesion was characterized using brightfield staining. Images of human postmortem tissue from the dorsolateral frontal cortex were obtained from the Boston University Alzheimer’s Disease Research Center (BU ADRC): 5 cases from 3 females and 2 males aged 75–105 with either a non-AD or low NIA-Reagan score, and with no other comorbid pathology were taken for the control group.

#### Human iPSC-derived microglia

A trisomic line (WC-24-02-DS-M) and an isogenic euploid control cell line (WC-24-02-DS-B) between passages 18 and 27 were generated from a female 25-year-old with Down syndrome and used in this study. The cell lines were produced, validated, and kindly provided by Dr. Anita Bhattacharyya from the University of Wisconsin-Madison. IPSCs were routinely passaged and cultured on Matrigel® (cat. 354277 Corning®, Corning, NY, USA) using mTeSR™ plus (cat. 85850, StemCell Technologies, Vancouver, BC, Canada). To generate iPSC derived microglia, iPSCs were differentiated into CD-43 expressing hematopoietic progenitor cells (HPCs) using the STEMdiff^TM^ Hematopoietic Kit (cat. 05310, STEMCELL Technologies) for 12 days. The expression of CD43, CD45, and CD34 was validated by flow cytometry. Cells expressing >90% CD43^+^ and >20% of CD34^+^/CD45^+^ on day 12 were confirmed to be HPCs, cryoprotected, and used for the subsequent differentiation into microglia. The microglial differentiation was performed on thawed HPCs (200,000 cells per well of a 6-well plate) using the STEMdiff^TM^ Microglia Culture System (cat. 100-0019 and 100-0020, STEMCELL Technologies) for 24 days according to manufacturer’s directions. On day 24, expression of >80% CD45^+^ and CD11b^+^ microglial markers was confirmed by flow cytometry to ensure proper differentiation. Cells were plated on poly-d-lysine coated glass coverslips (cat. A3890401, ThermoFisher, Waltham, MA, USA) at 100,000 cells/well and matured until day 30. Cells were fixed with 4% PFA for 30 min, washed twice with PBS, and stored at 4°C until use.

#### Primary neuronal culture

Primary neuronal cultures were prepared from postnatal day 0 (P0) mouse pups. At P0, brain tissues from WT and Ube3a 2x Tg male pups were dissected and digested, and then plated onto circular coverslips for neuron culture.

#### Drosophila

Fly strains *alrm-GAL4* (#67032) and *UAS-IVS-GFP* (#32201) were obtained from the Bloomington *Drosophila* Stock Center, crossed at 25°C, and raised at 29°C on standard cornmeal and molasses food. Flies aged to 2 days were dissected for immunohistochemistry. Only male flies were included in the experiment.

#### Sea anemone

*Nematostella vectensis* (*Nv*) were obtained from Maryland by Mark Martindale and Matt Gibson. The anemones were kept at ⅓ strength artificial seawater (ASW, ∼12 parts per 1000) in a dark incubator at 19°C with weekly water changes. Clonal pairs of *Nv* were generated by bisection and regeneration and then fed or starved for 30 days. Adult animals were bisected perpendicularly to the oral-aboral axis. Halves were placed in wells of a 24-well plate. Anemones regenerated for 30 days with feeding paused until tentacles formed from the aboral end. Adult *Nv* were fed freshly hatched brine shrimp (*Artemia*; brineshrimpdirect.com) and young polyps were fed ground *Artemia* in ⅓ ASW three times a week. Animals that were regenerated from aboral and oral ends were included. 40-day-old polyps that were fed or starved for 30 days were used.

#### Xenopus

Mucociliary epithelial MCCs of *Xenopus* embryos were used to study cilia formation. MCCs form >100 basal bodies and motile cilia that are important for generating directional fluid flow to clear pathogens from the airways or the tadpole skin. *Centrin4-cfp* was injected into embryos to identify targeted cells and basal bodies of MCCs. Knockdown of *cp110* in embryos was achieved with morpholino oligonucleotide (MO) injection to study the role of Cp110 in MCC ciliogenesis. To rescue cilia formation, *cp110* DNA was co-injected with *cp110* MO.

#### Corals

*Siderastrea siderea* (massive starlet coral) is one of the primary reef-building corals in Bocas del Toro, Panamá^32,33^. Previous work identified three cryptic lineages of *S. siderea* that occupy sites spanning the inshore reefs of Bocas del Toro^34^.

### Immunohistochemistry

#### Mouse

Sagittal sections of brains, hearts, and kidneys and sections of livers, lungs and spleens were captured at 16-μm thickness on Superfrost^TM^ slides and stored at -20°C until use. Tissue sections were rehydrated using PBS and permeabilized in blocking solution containing 0.5% Triton-X-100 (Sigma Aldrich, USA) and 5% bovine serum albumin (BSA) (Sigma Aldrich, USA) in PBS for 1 h at room temperature (RT). Tissues were incubated overnight at 4°C with rabbit anti- IBA1 antibody (1:500; Wako 019-19741) diluted in PBS containing 0.1% Triton-X-100 (Sigma Aldrich, USA) and 5% BSA (Sigma Aldrich, USA). After 5 10-min rinses in PBST, tissues were incubated with Alexa Fluor 647-conjugated donkey anti-rabbit secondary antibody (1:1000; Thermofisher Scientific A-31573) at RT for 2 h. After 5 10-min rinses in PBST, tissues were coverslipped with Prolong Gold Antifade Mountant containing 4′,6-diamidino-2-phenylindole (DAPI) (Cell Signaling, 8961).

#### Human

PLP was used to identify the areas of MS lesion and CD68 was used to identify the active rim of microglial cells (**Table 2**). For fluorescent staining, the formalin-fixed paraffin embedded (FFPE) block was sectioned at 7-µm thickness onto a slide, deparaffinized in xylene and rehydrated via an ethanol series. The section was subjected to antigen retrieval using citrate buffer at pH 6.0 followed by permeabilization using 0.5% Triton X-100 in Tris-buffered saline (TBS). After washing, the section was quenched in 0.3% H2O2 in TBS and washed again. Tissue was blocked for 1 h using 10% normal goat serum (NGS) and 10% Fc receptor inhibitor (Invitrogen, 14-9161-73) in TBS. Primary antibody for IBA-1 (Synaptic Systems, #234004) was applied with 10% NGS and incubated at 4°C overnight. After several washes in TBST, Alexa- Fluor 488-conjugated secondary antibody (Invitrogen, #A1073) was applied at RT for 3 h. After several washes, Sudan Black (MP Biomedicals, 152088) was applied to reduce autofluorescence, followed by washes and Hoechst Counterstain (Life Technologies, H357). Slices were coverslipped with 50% glycerol in TBS. Paraffin-embedded brain tissues from the BU ADRC were cut at 20-µm thickness. DAB (3, 3’-diaminobenzidine) immunohistochemical staining for IBA-1 (1:500, Wako, 019-19741) and nuclear counterstaining with hematoxylin was performed.

#### Human iPSC-derived microglia

Glass coverslips were treated with blocking solution containing 5% normal donkey serum in PBST (0.1% Triton X-100) at RT for 1 h. Cells on coverslips were incubated overnight at 4°C in rabbit anti-IBA1 (1:500; Wako, 019-19741) diluted in blocking solution. Coverslips were washed 3X for 10 min in PBST and incubated at RT for 1 h with Alexa- Fluor 546-conjugated donkey anti-rabbit antibody (1:1000; Invitrogen, A10040) in blocking solution. Coverslips were washed 3X 10 min in PBST. ProLong Gold Antifade Mountant with DAPI (Invitrogen, P36931) was applied to adhere coverslips to slides. Slides were stored at 4°C and light-protected until imaging.

#### Primary neuronal culture

Neurons plated on coverslips were fixed at day *in vitro* 8 (DIV8) in 4% PFA for 8 min and washed twice in 1X PBS. Cells were permeabilized for 10 min in 0.3% Triton X-100 (Sigma) in 1X PBS, followed by 2 washes in 1X PBS. Cells were blocked for 1 h in 10% NGS in 1X PBS. To label dendritic structures, cells were incubated overnight at 4°C in 1X PBS containing MAP2 monoclonal antibody (1:300; ThermoFisher, #13-1500). The next day, cells were washed 3X with cold 1X PBS, followed by incubation in 1X PBS containing Alexa Fluor 488 or 555-conjugated mouse IgG secondary antibody (1:500; Invitrogen, #A-21422. Cells were counterstained with Hoechst (1:10,000 in 1X PBS; ThermoFisher, H3570) for 30 sec followed by 2 washes in 1X PBS. Coverslips were mounted onto slides with Prolong Gold (ThermoFisher, P36930) and left to dry in the dark overnight before imaging.

#### Drosophila

Male flies were fixed in 4% PFA diluted in 1X PBS and 0.5% Triton X-100 (PBT) at RT for 3 h. Flies were washed 4X in PBT for 15 min each with rotation and dissected in PBS. Whole brains were blocked at RT for 1 h in PBT containing 0.5% BSA and 5% NGS (PBANG) and then incubated in PBANG containing rabbit anti-GFP (1:100; Torry Pines Biolabs, TP401) at 4°C for 2 nights. Brains were washed 4X in PBT at RT and incubated in PBANG containing goat anti-rabbit Cy3 (1:200; Jackson ImmunoResearch, 111-165-003) for 2 nights at 4°C. Brains were washed 3X in PBT. Samples were placed in VectaShield with DAPI (Vector Labs) overnight at 4°C and subsequently mounted on slides.

#### Sea anemone

Whole-mount immunofluorescence staining of NF-κB was performed using a custom antibody and Texas Red-labeled secondary antibody to visualize NF-κB levels in *Nv*. For immunofluorescence, *Nv* polyps were fixed in 4% formaldehyde in ⅓ ASW overnight at 4°C and then washed 3X with 0.1% Triton X-100 in PBS (PTx). Polyps were then microwaved in warm urea (5% w/v) at the lowest setting for 5 min, cooled at RT for 20 min, then washed 3X with PTx. Polyps were blocked overnight in PTx containing 5% NGS and 1% BSA at 4°C on a nutator. Anti-Nv-NF-κB antibody diluted 1:100 in blocking buffer was used overnight at 4°C. Polyps were washed 4X with PTx and incubated in Texas-red-conjugated anti-rabbit secondary antibody (1:160; Invitrogen, #T-2767). DAPI was added to a final concentration of 5 mg/ml to visualize nuclei.

#### Xenopus

Whole embryos were fixed at embryonic stages 30-32 overnight at 4°C, washed 3X 15 min with PBS, 2X 30 min in PBST (0.1% Triton X-100 in PBS), and blocked at RT in PBST-CAS (CAS blocking; ThermoFisher #00-8120) for 1 h. To visualize cilia, embryos were incubated overnight at 4°C in mouse monoclonal anti-acetylated-α-tubulin antibody (1:700; Sigma, #T6793) and Alexa Fluor 555-labeled goat anti-mouse antibody (1:250; Molecular Probes, #A21422) in CAS Blocking.

### Image acquisition and processing

#### Mouse

Z stacks of whole organ sections were acquired on a ZEISS AxioScan 7 slide scanner using a 20X 1 NA objective lens at 1X zoom and 1.1 binning. Images were sampled with a voxel size of 0.325-μm X 0.325-μm X 1-μm for the DAPI (excitation wavelength 353-nm; emission wavelength 465-nm) and Alexa Fluor 647 (excitation wavelength 650-nm; emission wavelength 665-nm) channels. Image tiles were stitched, processed for maximum intensity projections (MIPs), cropped to obtain >200 cells per image, and exported as TIFF images using the ZEISS ZEN pro software (ZEN 3.2 blue edition). At least 4 images were analyzed per organ/mouse.

#### Human

Tissue sections from MS patients were scanned on a Leica Microsystems DM6B microscope a 40X objective lens at 0.329-μm per pixel. Each tile was focused using the “Auto- focus” function and subsequently captured in multiple z-stack images of 6-µm thickness in total. Z-interval was set depending on tissue quality. Tiled images were merged with the “Mosaic merge” function to include a 20% overlay area. The “Projection” function was used to obtain MIPs that contained the best signal intensity for each pixel. Images were cleaned up using “Instant Computational Cleaning” under the “Thunder” function before they were exported in TIFF for image processing in ImageJ. Images containing MS lesions were characterized by PLP, CD68, and IBA1 expression and precisely cropped for morphometric analysis. Tissue sections from BU ADRC were imaged using an Aperio GT450 slide scanner at a 40X magnification. Images were processed using the “extract region tool” in the Aperio ImageScope Pathology Slide Viewing Software to generate 10–12 cropped images measuring 1000-µm X 1000-µm each from the white and gray matter. To enhance the DAB signal and minimize the noise contributed by the nuclear counterstain, each duplicated MIP was thresholded to exclude hematoxylin and subtracted from the original MIP using “Image Calculator” to generate a new 8-bit TIFF image for AutoMorFi analysis.

#### Human iPSC-derived microglia

Images were acquired on an Olympus Fluoview FV300 using a 40X oil immersion objective lens and the FV31S-SW Viewer software. Images were captured in DAPI (excitation wavelength 353-nm; emission wavelength 465-nm) and Alexa-Fluor 546 (excitation wavelength 556-nm; emission wavelength 573-nm) channels.

#### Primary neuronal culture

Neuronal images were captured using a Carl Zeiss Axiovert inverted fluorescence microscope with a 40X 1.3 NA oil immersion objective lens and the AxioVision Release 4.5 software.

#### Drosophila

Images of the central brain were captured on a Nikon C2+Si confocal microscope using a 40X 1.25 NA water immersion objective lens and NIS-Elements software. Fifty planes 1- μm apart were captured for the Z stacks of each brain with a resolution of 1024 X 1024 pixels starting from the surface of the central brain at the first point where the antennal lobe becomes visible. MIPs were generated in ImageJ at serial steps of 10 (e.g., 1-10, 11-20, …, and 41-50) to distinguish individual glial cells in each brain.

#### Sea anemone

Images were acquired on a Nikon C2 Si confocal microscope and processed using ImageJ.

#### Xenopus

Images were acquired using a ZEISS LSM700 confocal microscope. ImageJ was used for Z-stack analysis and processing.

#### Corals

Photos of representative coral colonies of each of the three cryptic lineages were taken from three sites across Bocas del Toro (Hospital Point, Cristobal Island, and STRI Point) using an Olympus Tough TG-6 camera on the underwater macro setting. For each individual coral (*N =* 52), macro photos focused on individual corallites were taken in triplicate, with each of the three photos taken on a different location on the coral surface. Photographs were converted to 8-bit binary images and scaled using the 2-mm size standard with the Set Scale function in ImageJ/Fiji. Images were individually adjusted by 0.35% contrast enhancement, smoothing, thresholding, and despeckling.

#### AutoMorFi morphometric analysis of single cells

Images were blinded before the individuals who did not generate the primary data ran the AutoMorFi macro except for iPSC-derived microglial cells and *Drosophila* astrocyte-like glia which were known to the persons who performed image analysis. A different set of individuals performed quality control to validate the AutoMorFi output. They compared the outlines of the detected “soma” and “whole cell” with cells in the original images to assess for false positive or false negative detections and inaccurate object segmentation in consultation with authors of the original experiments. Incorrectly detected cells or objects were removed from the .csv output files. Output images with missing cells were reanalyzed using improved diameter measurements. Procedures for running the AutoMorFi macro in ImageJ/Fiji are detailed in **Supplementary Notes 1–4**. Each single-cell measurement of “soma”, “whole cell”, and “skeletonized” structure (generated by ImageJ/Fiji from the whole-cell outline) derived from AutoMorFi were matched by the coordinates of the objects using the Python packages numpy, pandas, and sklearn.neighbors, and collated in a single .csv data file (**Supplementary Note 6**). AutoMorFi generated 33 morphometric measurements that were in part used in arithmetic functions that produced additional parameters resulting in a total of 47 quantitative characteristics per cell (**Table 1**).

#### AutoMorFi morphometric analysis of single objects

Image blinding and data validation were carried out as described for AutoMorFi analysis of single cells. To customize AutoMorFi for measuring non-cellular objects, the use of the pipeline was modified as follows.

#### Sea anemone

The diameters of several whole polyps were measured from the DNA channel to obtain a range of representative values. While executing AutoMorFi, the Analyze Particle plugin was set to “Overlay” instead of “Mask” to save the polyp outline as an overlay that was now considered the “whole cell”. The overlay was renamed and manually saved as a TIFF image in the output folder before continuing with the AutoMorFi macro. A region of interest (ROI) was marked in the background of the image and added to the ROI manager to generate a second pseudo-object to allow the acquisition of all standard parameters of the whole single polyp (see Error 7 in **Supplementary Note 3**). The macro was continued until the end. Measurements from the marked ROI and “soma” were excluded from the data analysis. “Whole cell” and “skeletonized” objects based on whole-polyp outlines contributing to 21 measured parameters were matched by their coordinates and collated in a single .csv data file. To acquire the total fluorescence intensity measurement of DNA and NF-κB signals, the overlay outline or “whole cell outline” that was saved earlier was added to the ROI manager in ImageJ/Fiji for each individual channel per organism.

The measured total fluorescence intensity values were normalized to the polyp area within the whole cell outline. The NF-κB area fraction represents the percentage of non-zero pixels within the polyp outline. An additional 12 calculated parameters (**Table 1**: 34, 35, 38–42, 46, 47; total DNA fluorescence intensity, total NF-κB fluorescence intensity, and NF-κB area fraction) resulted in a total of 33 quantitative characteristics per polyp. Area fraction (%Area) is acquired from the table of results generated from ImageJ/Fiji (Analyze > Measure).

#### Xenopus

Focusing only on the acetylated-α-tubulin channel of the imaged MCCs, diameters of several representative objects indicated by acetylated-α-tubulin staining were measured across images to obtain a range of values for running the AutoMorFi macro. Measurements of “soma” were excluded. “Whole cell” and “skeletonized” outlines were matched by their coordinates. Total acetylated-α-tubulin fluorescence intensity was measured for each MCC. A total of 43 quantitative characteristics was obtained per MCC by including the calculated parameters (**Table 1**: 34, 35, 38, 39, 40, 41, 42, 46, 47), total acetylated-α-tubulin fluorescence intensity, and morphometric measurements of manually segmented MCCs based on *centrin4-cfp* signals (**Table 1**: 1 to 12). **Corals.** Representative diameters of several corallites were acquired across coral lineages. AutoMorFi adapted for color photography (**Supplementary Note 5**) was used. Measurements of incomplete corallites, corallites covered by sand, and “somas” were excluded. “Whole cell” and “skeletonized” parameters contributed to 21 morphometric measurements. Additional calculated parameters (**Table 1**: 34, 35, 38, 39, 40, 41, 42, 46, and 47) resulted in a total of 30 quantitative characteristics per corallite.

### Data visualization

Unpaired t-tests, pairwise comparisons of morphometric parameters, and principal component analysis (PCA) were performed between or across groups to determine the need for dimensionality reduction^5,35^ using tidyverse, ggplot2, and heatmap R packages. Redundant morphometric features with high correlation coefficients between 0.5 and 1 were removed to mitigate bias in subsequent cluster analysis. Unsupervised K-means clustering analysis was performed on the reduced datasets using the R factoextra package. For data visualization, UMAPs were generated using tidyverse, dplyr, ggplot2, and umap R packages, and heatmaps were generated using dplyr, ggplot2, and heatmap R packages in R.4.3.2.

### Statistical analysis

Paired and unpaired t-tests were performed using GraphPad Prism10. Data are presented as mean ± s.e.m. and considered statistically significant at *p* < 0.05.

## SUPPLEMENTARY NOTE 1

### Prerequisites to running AutoMorFi

**Step I:** Download and add the following plugins to the “plugins” folder of your ImageJ:

- “Excel.macro.extensions-3.3.2-jar-with-dependencies.jar”
- “script-editor-0.7.4-SNAPSHOT.jar”

To find the “plugins” folder on Mac:

- Go to Finder > Desktop > Search: “Fiji”
- Right-click on the Fiji logo > “Show package contents”
- Open the “plugins” folder
- Copy-and-paste the downloaded plugins

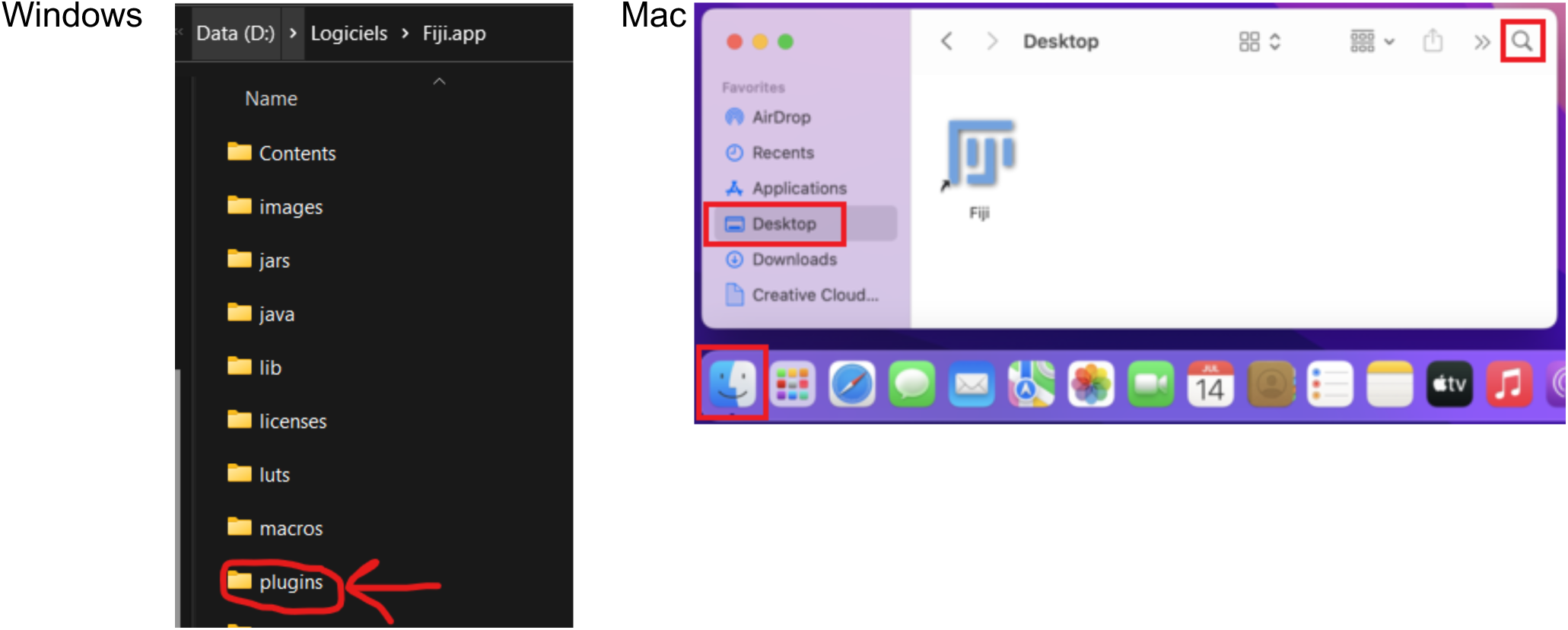

**Step II:** Update ImageJ/Fiji

- Start ImageJ/Fiji and select Help > Update… > Manage update site
- Select: 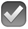 Excel Functions 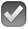 Neuroanatomy
- Unselect: □ResultToExcel
- Close the “Manage update sites” window
- Close the “ImageJ Updater” window
- Restart ImageJ/Fiji

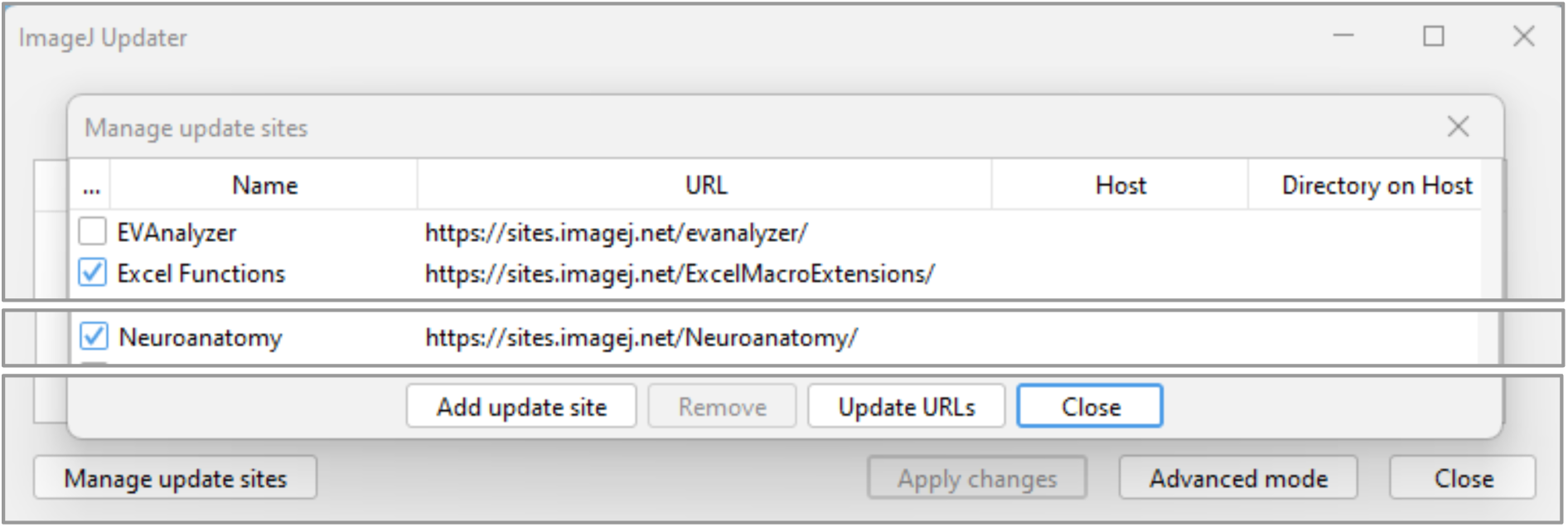

**Step III:** Range of measurements

1. Ensure your version of Fiji/ImageJ is updated before each analysis.
2. Set your Fiji/ImageJ as follows:

- Analyze > Set measurements
- Select the top 18 criteria
- Click “OK”

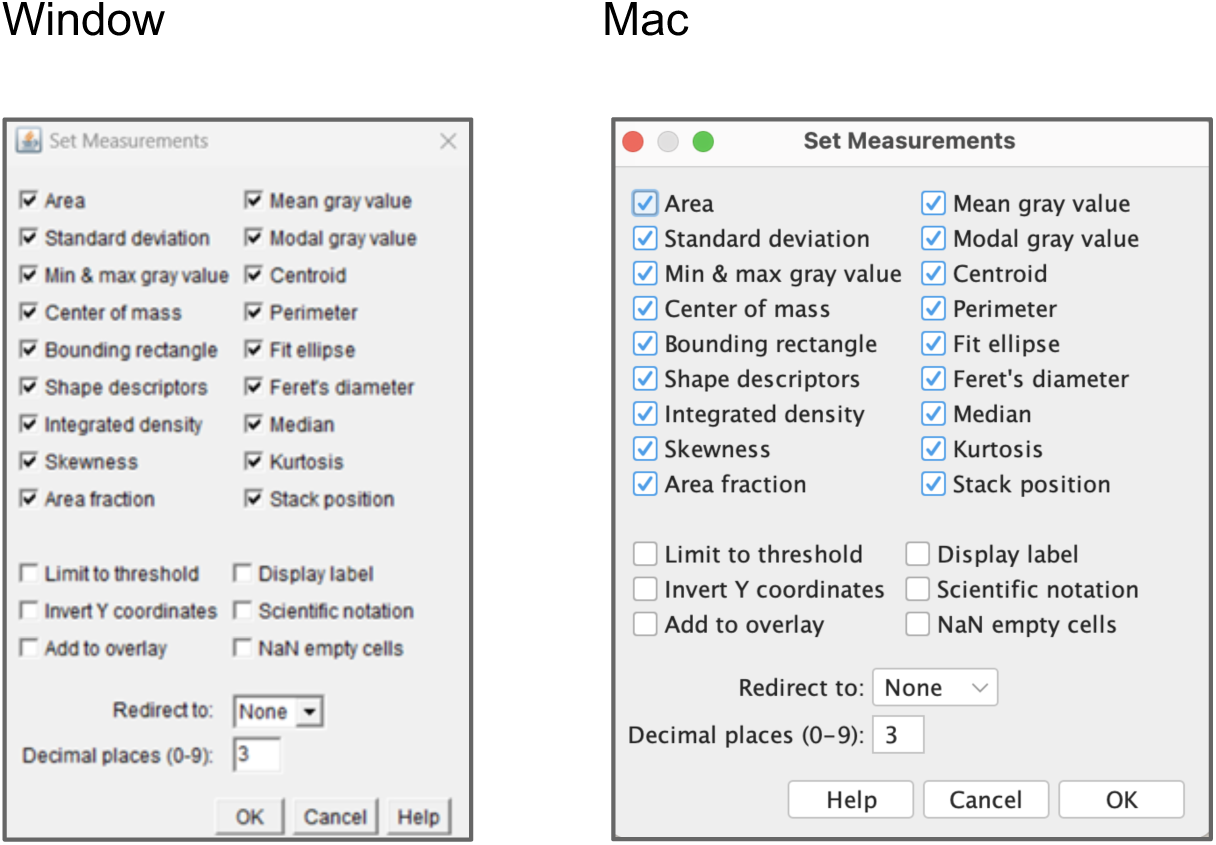

3. AutoMorFi requires the user to enter the size range of the cell soma and whole cell areas. To get these values:

- Drag and drop the image you want to analyze onto Fiji/ImageJ
- Make sure to work with a single-channel .tif image that is 8-bit and scaled (i.e., unit per pixel).
- Use the ‘straight’ tool to measure the diameter (in microns) of 5–10 cell somas and whole cells representing the average, smallest, and largest objects in the image.
- With these diameter measurements, estimate the size ranges needed in the Analyze Particles wizard embedded within AutoMorFi. For example, if the soma diameters measured are 6.74, 7.56, 8.43, 13.74, and 12.33, the lower threshold value would be obtained as 6.74^2 = 45.43 and the upper threshold value would be obtained as 13.74^2 = 188.79. The range entered in the wizard could be 42- 200.
- Optimization tip: the ranges of soma and whole cell areas depend on the type of image and objects you are working with (e.g., cell type, cell density, method of image acquisition, etc.). After the first run, evaluate the accuracy of your soma and whole cell ranges by looking at the soma (SomaOutlines.tiff) and whole cell (WholeCellOutlines.tiff) outlines in your output folder.

- too many nonspecific objects identified –> increase lower threshold value
- small objects not captured –> decrease the lower threshold value
- big objects not captured –> increase the upper threshold value

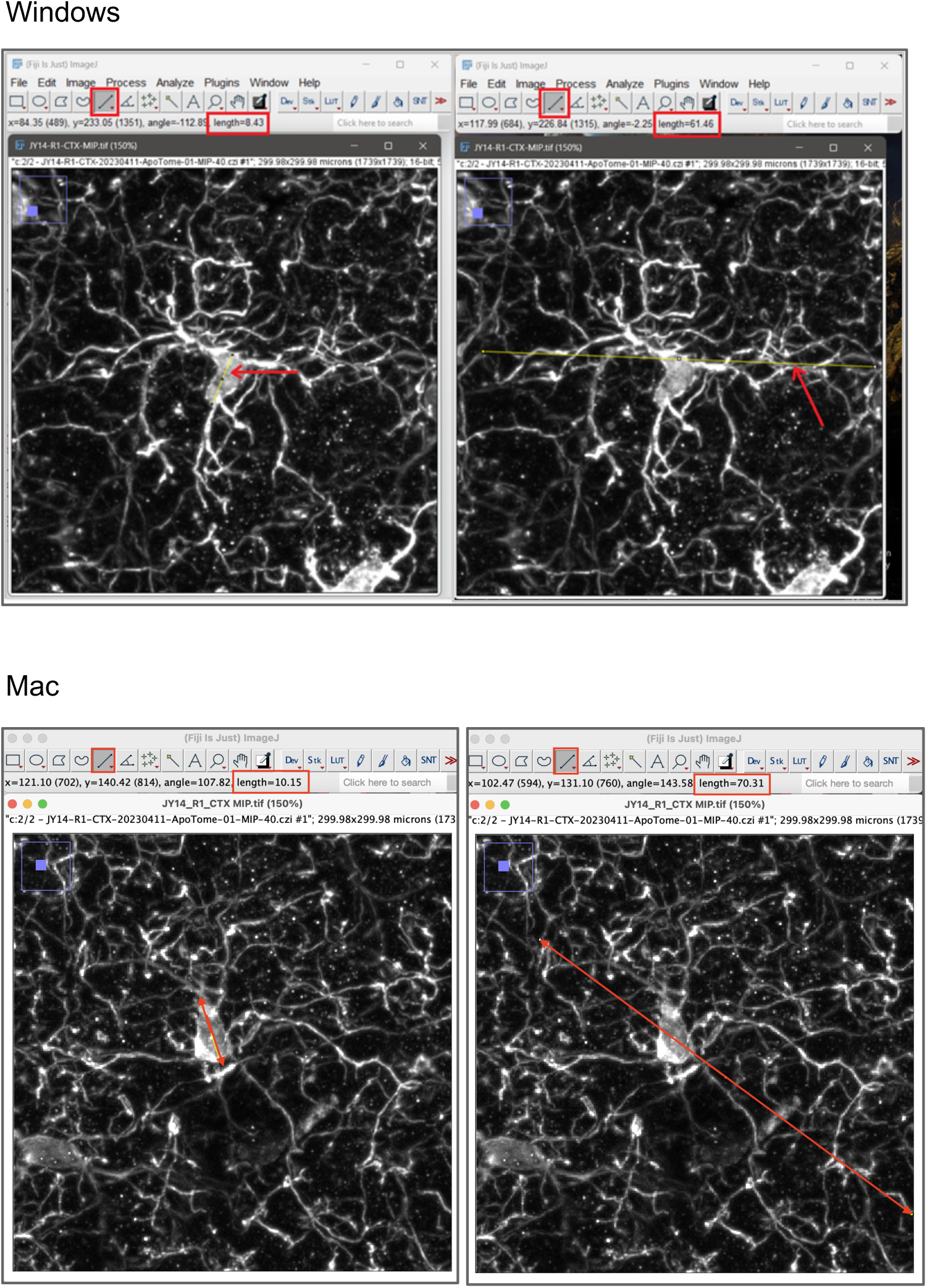

## SUPPLEMENTARY NOTE 2

### Running the AutoMorFi macro

AutoMorFi analyzes **one scaled single-channel 8-bit TIFF image** per cycle. Make sure the folder path leading to your image contains **<u>no space, slash, or other special characters</u>**. Use only letters, numerals, underscore, and dash.

Close any window that is open on Fiji/ImageJ. To start the AutoMorFi macro (AutoMorFi_3.ijm):

1. drag and drop the macro file in Fiji/ImageJ > press [Run]
2. select your image (“xxx.tif”) > Click “Open”

#### Step 1: object identification by thresholding

On your screen, the original image (named “Temp”) and the “Skeleton” image are displayed. In the “Threshold” window, adjust the threshold of the “Skeleton” image:

1. select the “Skeleton” image (an error message will result if a different image is selected)
2. adjust the auto threshold method (ex: Default)
3. Tip: set the display method to “Red” (instead of B&W) in the “Threshold” window
4. adjust the threshold using the first sliding bar
5. **<u>click “Apply”</u>** in the “Threshold” window
6. click “OK” to continue the analysis

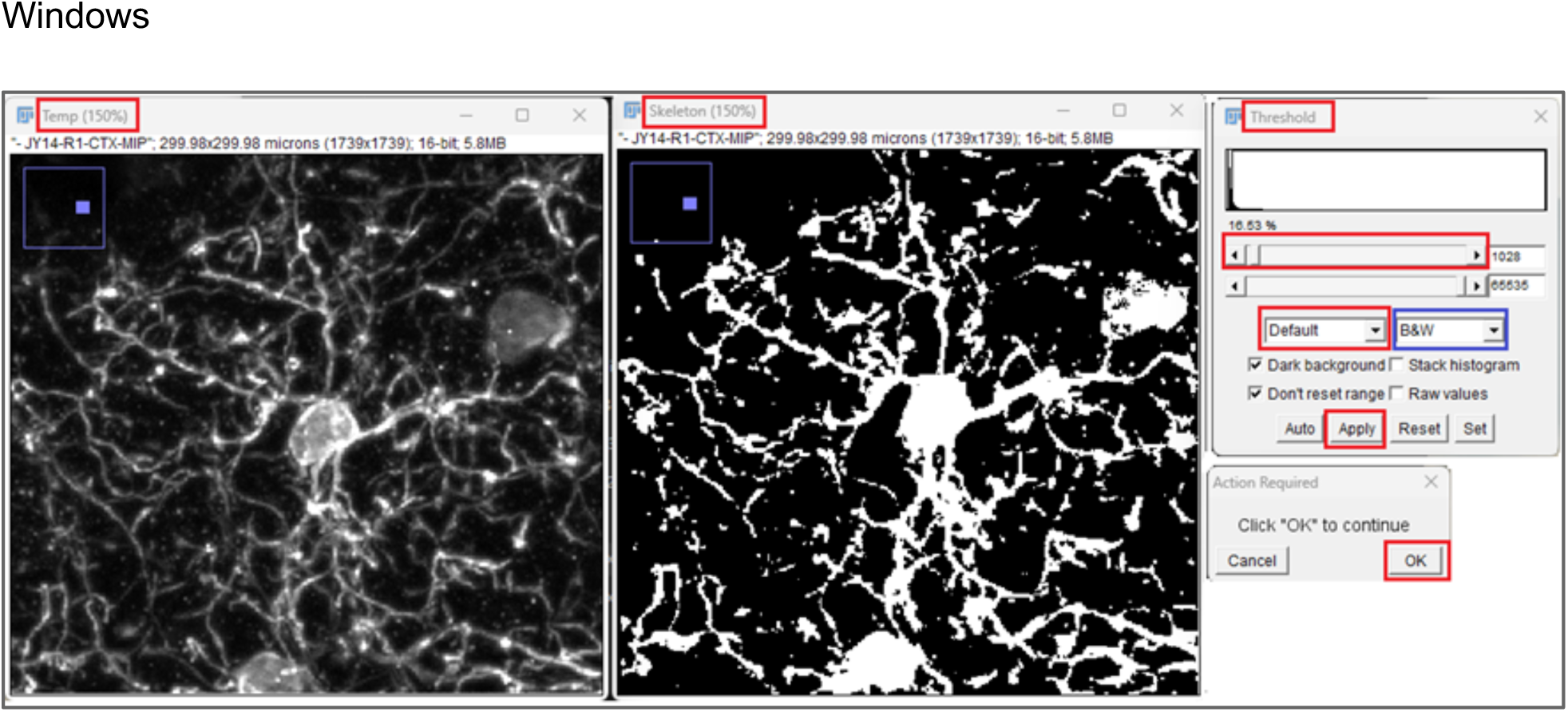
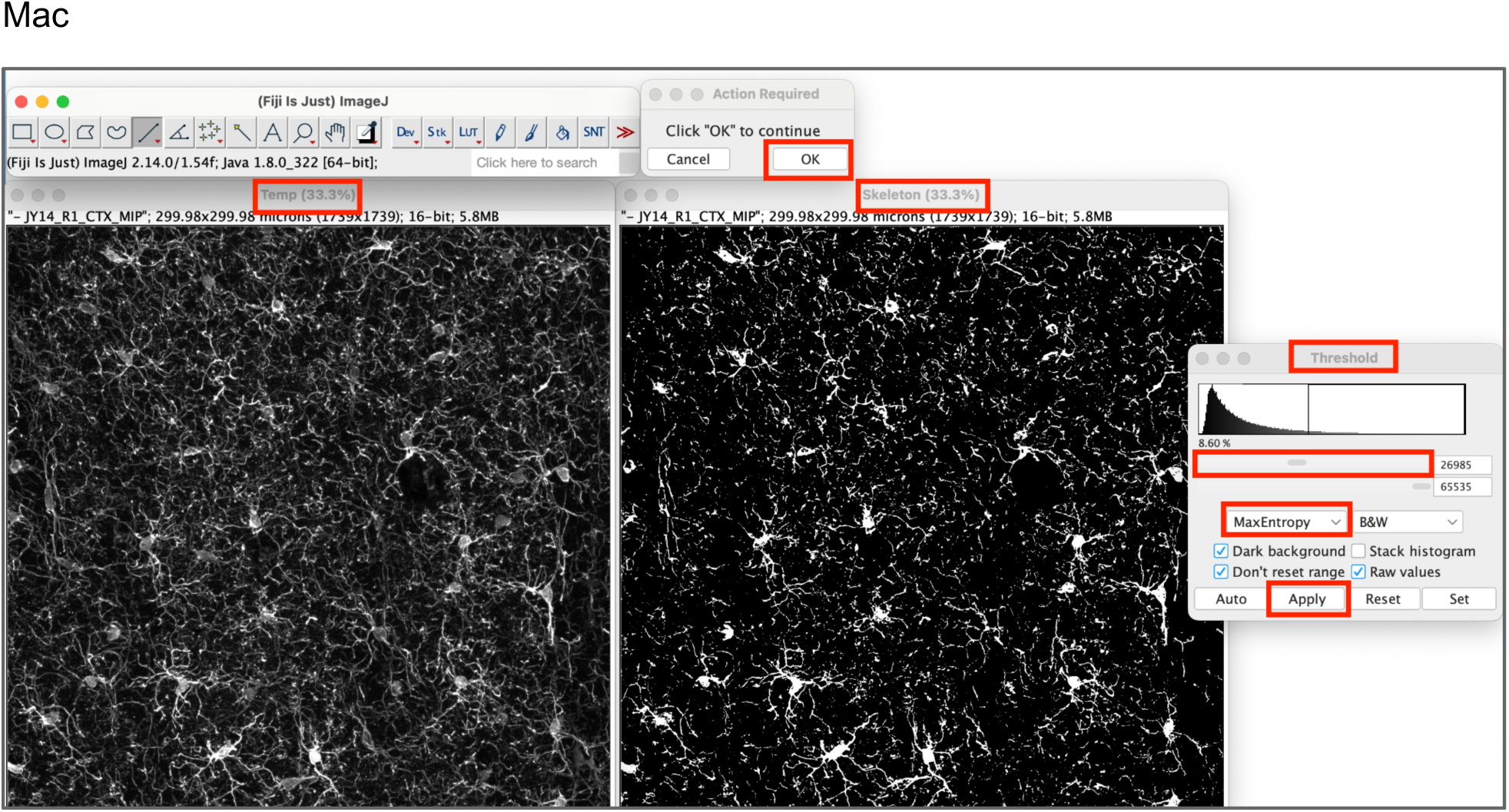

#### Step 2: connect disrupted processes using dilation (for objects with branches)

- Dilate Option window: if your cells are not branched, select “No” > click OK
- Dilate Option window: to connect disrupted processes, select “Yes” > click “OK”
- to **finish** the dilation: select “Finished” > “OK”
- to **revert** the dilation: select “Revert” > “OK” (only one dilation can be reversed)
- to **continue** dilating: select “Continue” > “OK” > “Yes” > “OK” Windows Mac

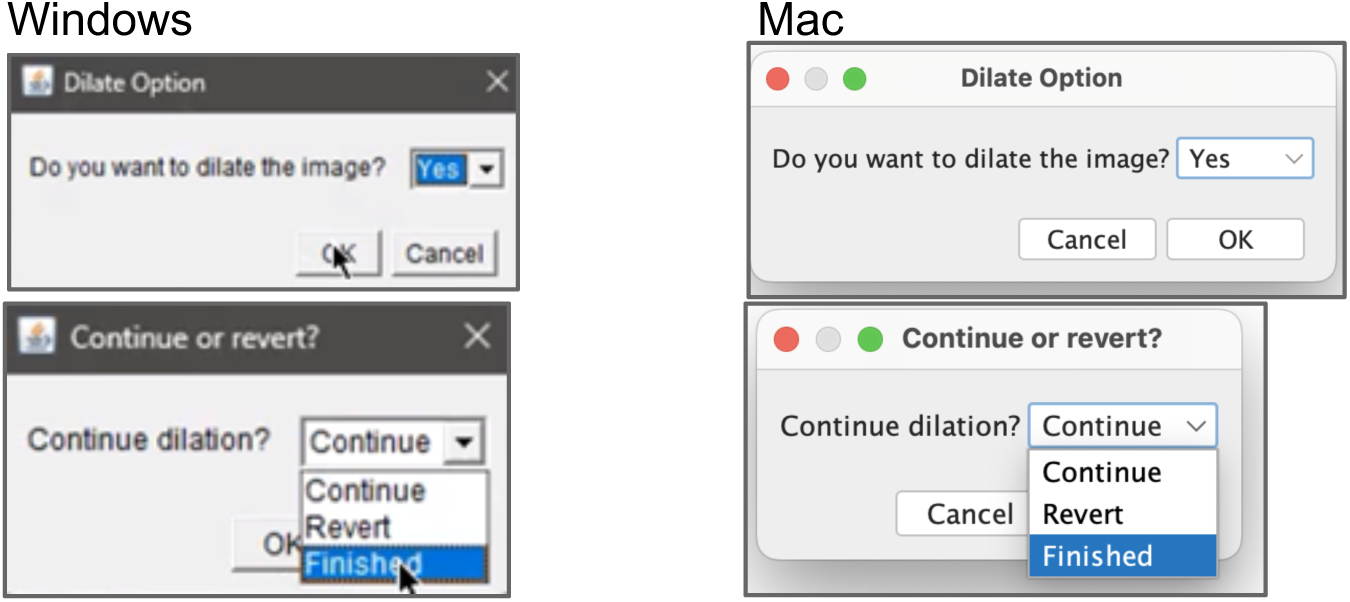

#### Step 3: object detection by diameter ranges

1. The **first** Analyze Particles window will analyze your cell **soma**

a. In “Size (micron)^2”, enter the calculated soma size <u>range</u>
b. Show: <u>Masks</u>, 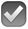 Display results, 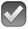 Add to manager, 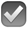 Overlay > “OK”
2. The **second** Analyze particles window will analyze your **whole cell**

In “Size (micron^2)”, enter the calculated whole cell size <u>range</u>
Show: <u>Masks</u>, 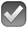 Display results, 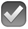 Add to manager, 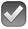 Overlay > “OK”

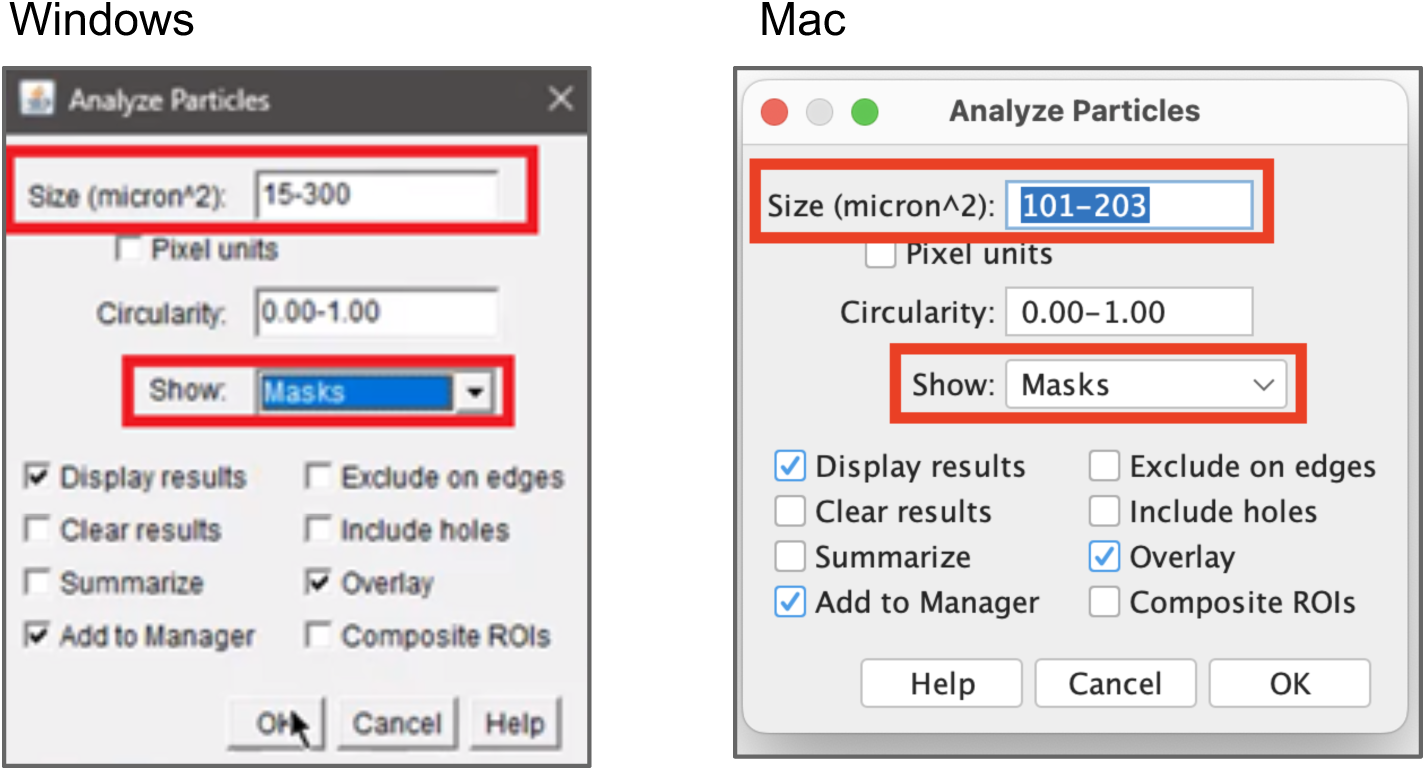

#### Step 4: deletion of false positive objects in the ROI manager

1. Select the objects by their number in the ROI manager (click on “Labels” if the numbers are not visible)
2. Click “Delete”
3. When you are done, click “Update” > “Deselect”
4. Click “OK” in the “ROI Deletion” window

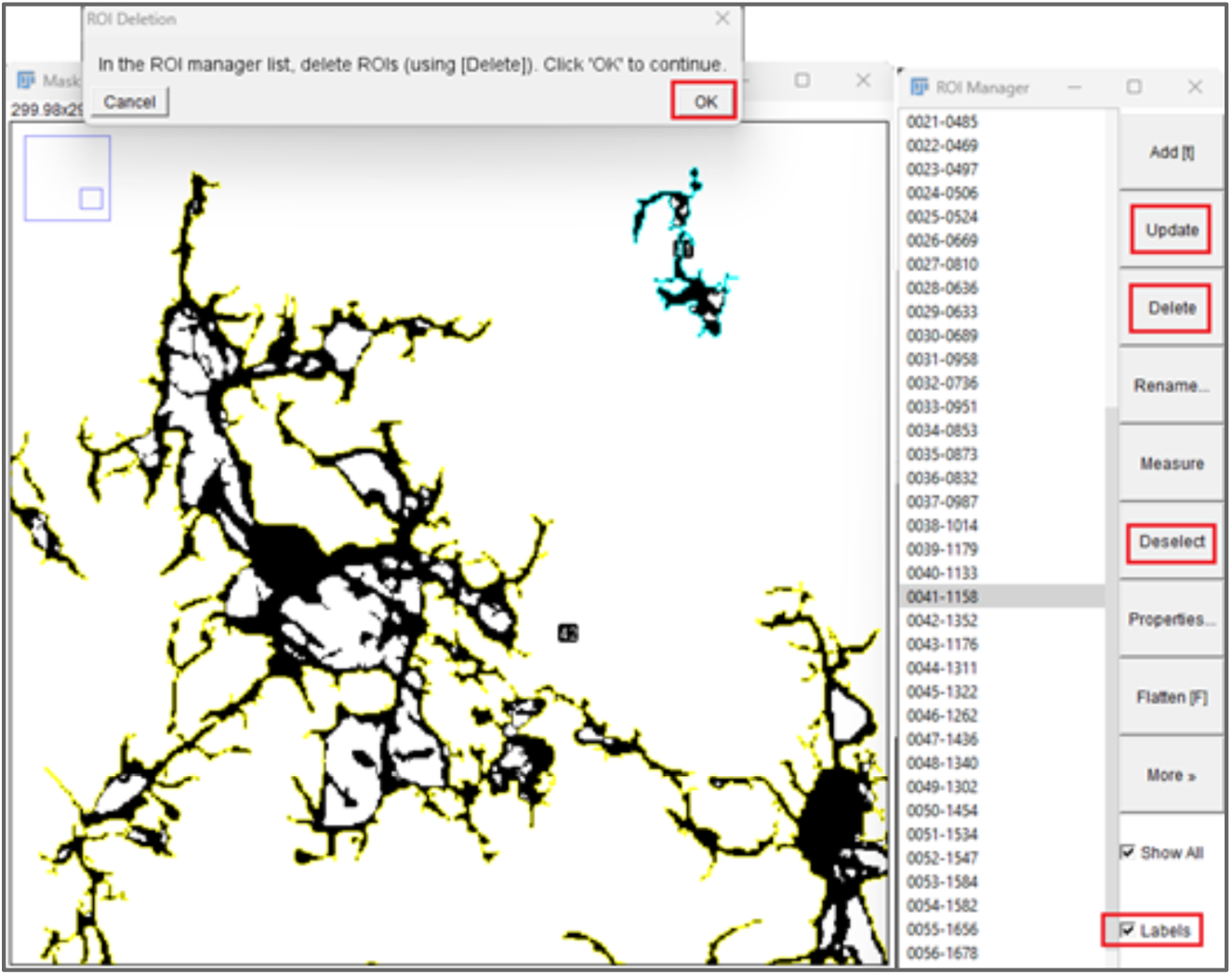

Let the macro run until the “Finish” window appears.

## SUPPLEMENTARY NOTE 3

### Troubleshooting for AutoMorFi

#### Error 1. Macro does not start

The image format, name, or image path is incorrect.

**Solution 1:** AutoMorFi only works on one-channel 8-bit TIFF images.

**Solution 2:** Make sure the folder path leading to your image contains no space, slash or special characters. Use only underscore or dash.

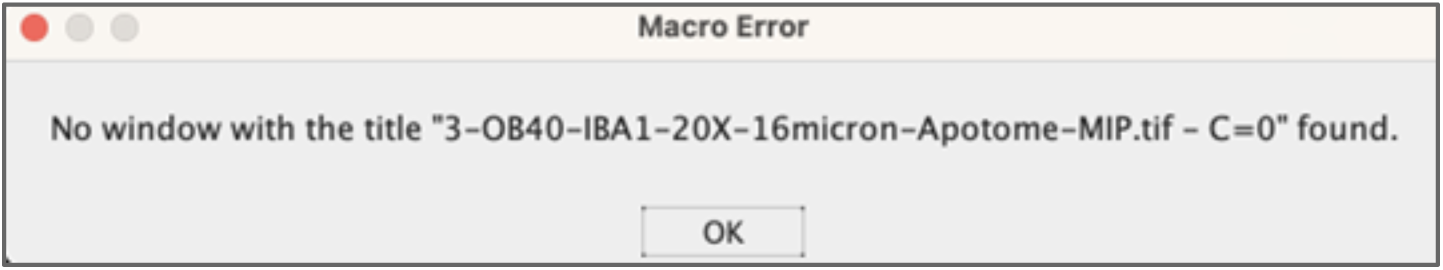

#### Error 2. Macro does not continue after step 1

A different image than “Skeleton” was selected for thresholding during step 2. The user did not click

“Apply” in the Threshold window before clicking “OK” in the Action Required window.

**Solution:** Make sure to adjust the threshold of the “Skeleton” image in the Threshold window and click “Apply” in the Threshold wizard before “OK” in the Action Required window.

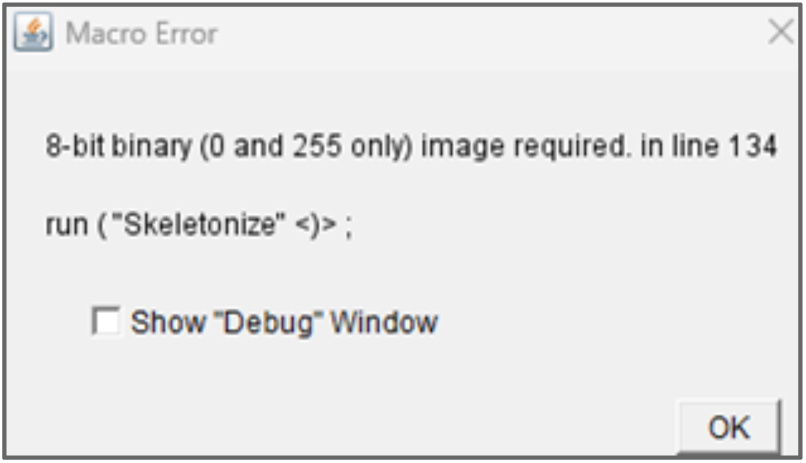

#### Error 3: Macro does not continue after step 3

No particle was detected based on the particle range set by the user in step 3 of AutoMorFi. The macro does not proceed without the “Results” window.

**Solution 1:** Check that the calculated size range provided for particle detection is correct (see **Step III** in **Supplementary Note 1**).

**Solution 2:** Locate the .tif image “Binary” in the output folder of the aborted AutoMorFi run. If the objects of interest are incorrectly represented in “Binary”, adjust the extend of thresholding in step 1 (**Supplementary Note 2**).

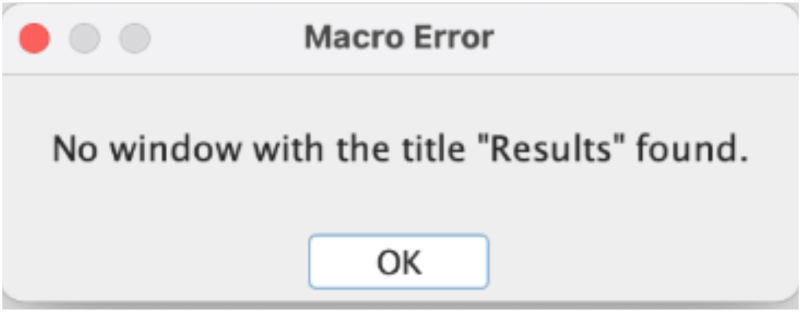

#### Error 4. Only one row of data is shown in the BranchInformation.csv and SkeletonResult.csv output files

The user did not click “Update” followed by “Deselect” in the ROI Manager window before clicking “OK” during **step 4 (deletion of false positive objects in the ROI manager).**

#### Error 5. Unresponsive macro or to interrupt the current run

**Solution:** Click “Kill”, at the bottom of the macro window.

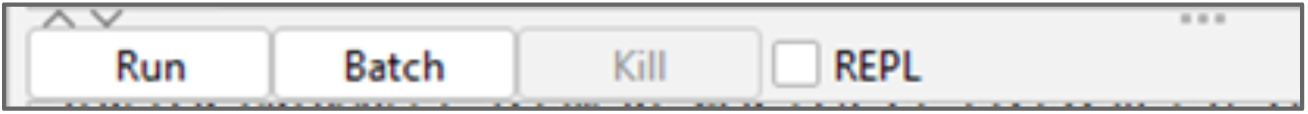

#### Error 6. “Default” thresholding incorrectly detects “soma” of the objects

AutoMorFi has a built-in automatic thresholding function for “soma” identification instead of the manual thresholding for whole-cell object identification in step 1 (i.e., object identification by thresholding). This can lead to incorrect “soma” identification in the SomaOutline.tif output.

**Solution:** Choose the appropriate AutoThreshold method to use before running AutoMorFi:

- Open your image on ImageJ/Fiji to identify the optimal AutoThreshold method

- Select: Image > Adjust > AutoThreshold (https://imagej.net/plugins/auto-threshold)

- In the AutoMorFi macro window, replace “Default” with the optimal AutoThreshold method (case-sensitive) in line 68 of the standard AutoMorFi code in **Supplementary Note 4**

- Run AutoMorFi as usual

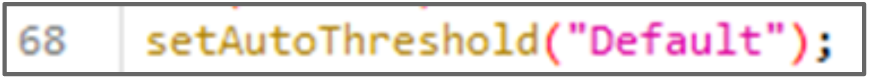

#### Error 7. Macro does not generate output from image containing only one object

AutoMorFi requires a minimum of 2 objects per image to run.

**Solution:** Create a second object in the ROI manager after the primary object during step 4:

- In the ROI manager, click “More” > “Draw”

- Draw a rectangle in the background of the image (i.e., not on the object of interest)

- Adding the second object in the bottom right corner of the image is recommended for coordinate-based synchronization of “soma”, “whole cell”, and “skeleton” outputs using the cell matching app in **Supplementary Note 6**

- When completed, click “Update” > “Deselect”

- Click “OK” in the “ROI Deletion” window

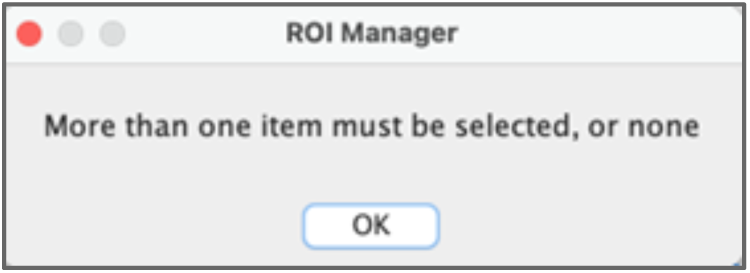
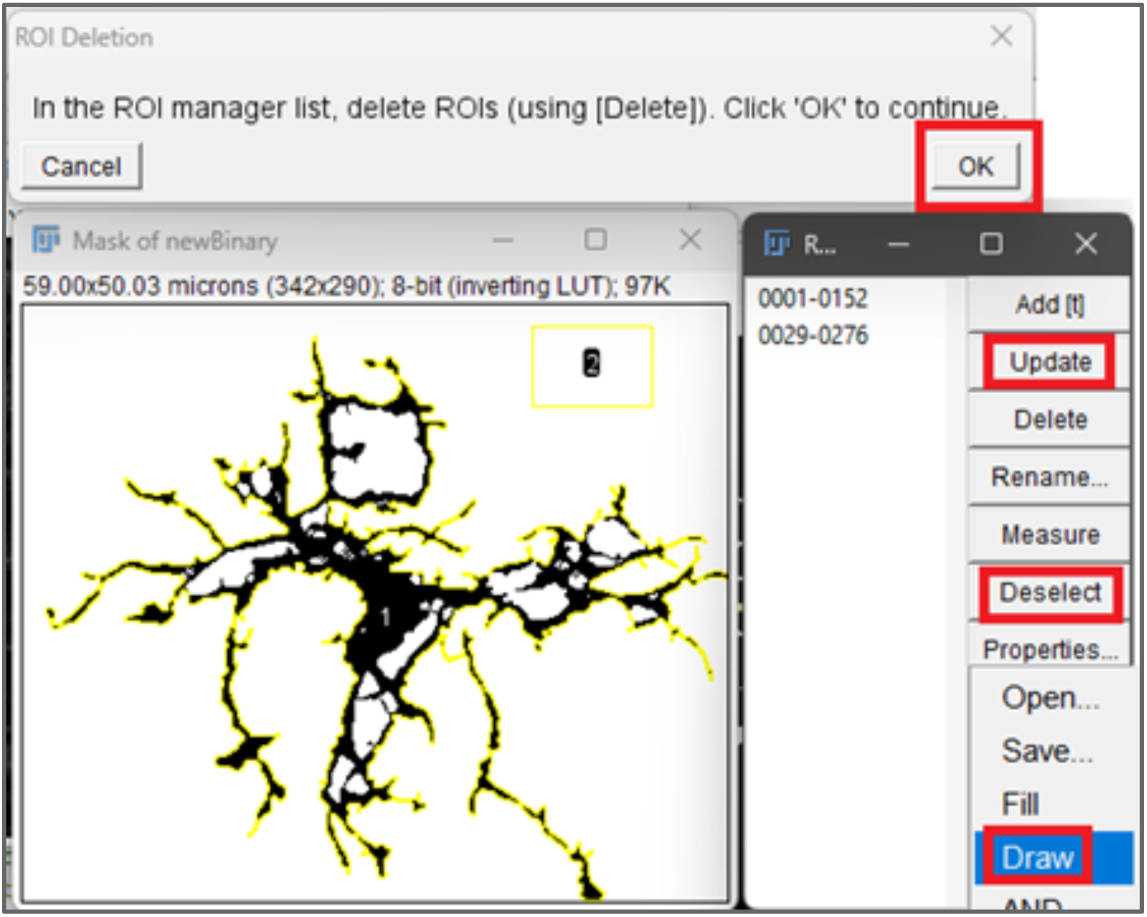

## SUPPLEMENTARY NOTE 4

### CODE FOR AUTOMORFI MACRO (IMAGEJ/FIJI-COMPATIBLE)

~~~
//AutoMorFi_3 macro
// clean up: makes sure no data is stored from previous macro use
// close all images close(“*”);
// empty the ROI manager roiManager(“reset”);
// empty the results table run(“Clear Results”);
// configure that binary image are black in background, objects are white setOption(“BlackBackground”, true);

//Obtain Image Path, Folder Path
path = File.openDialog(“Choose a File”); basename = File.nameWithoutExtension(); dirname = File.getParent(path);
folder = dirname + File.separator + basename;

//Create Result Folders
if (File.exists(folder)) {
 print(“Directory already exists, override the existing directory”);
 list = getFileList(folder);
 for (i = 0; i < list.length; i++) {
    File.delete(folder + File.separator + list[i]); // Delete each file
    }
}
else {
 File.makeDirectory(folder);
}

//Open Image thru Bio-Format Importer run(“Bio-Formats Importer”, “open=”+path);
imageOriginPath = folder + File.separator + basename; saveAs(“tiff”, imageOriginPath);

//Get Image size, Set scales
getPixelSize(unit, pixelWidth, pixelHeight);
scale = 1 / pixelWidth;
nameOrigin = getTitle(); microWidth = getWidth() / scale;
microHeight = getHeight() / scale;

//All Output Directories
binaryPath = folder + File.separator + “Binary”;
somaResultsPath = folder + File.separator + “SomaResults.csv”; somaOutlinesPath = folder + File.separator + “SomaOutlines”;
wholeCellResultsPath = folder + File.separator + “WholeCellResults.csv”;
wholeCellOutlinesPath = folder + File.separator + “WholeCellOutlines”; skeletonPath = folder + File.separator + “Skeleton”;
skeletonResultsPath = folder + File.separator + “SkeletonResults.csv”; branchInformationPath = folder + File.separator + “BranchInformation.csv”;

//Duplicate original input twice. One for skeleton, the other for threshold reference.
run(“Duplicate…”, “ “);
rename(“Skeleton”);
run(“Duplicate…”, “ “);
rename(“Temp”);

//Process Binary Image
selectWindow(basename+”.tif”);
run(“Enhance Contrast”, “saturated=0.35”);
run(“Apply LUT”);
run(“8-bit”); setAutoThreshold(“Default”);
run(“Convert to Mask”); run(“Invert LUT”);
run(“Make Binary”);
run(“Despeckle”);
rename(“Binary”);

//Process Skeleton, Manually set threshold
selectWindow(“Skeleton”);
run(“Threshold…”);//https://imagej.net/plugins/auto-threshold#available-methods waitForUser; //User adjust threshold here
close(“Threshold”);
close(“Temp”);

//Dilation Dialogs
selectWindow(“Skeleton”);
stack_count = 0; //Keep track of the number of dilations
rename(“Mask_” + stack_count);
while (true) {
  //Creating dialog with dilate option “Yes”/“No”
  Dialog.create(“Dilate Option”);
  Dialog.addChoice(“Do you want to dilate the image?”, newArray(“Yes”, “No”));
  Dialog.show();
  choice = Dialog.getChoice();

  //When choosing “Yes”, dilate the image from the previous stack
  if (choice == “Yes”) {
    selectWindow(“Mask_” + stack_count);
    run(“Duplicate…”, “title=Temp”);
    run(“Dilate”);
    stack_count++;
    selectWindow(“Temp”);
    rename(“Mask_” + stack_count);

    //Creating dialog with options “Continue”, “Revert”, “Exit” Dialog.create(“Continue or revert?”);
    Dialog.addChoice(“Continue dilation?”, newArray(“Continue”, “Revert”, “Finished”));
    Dialog.show();
    secondChoice = Dialog.getChoice();
    //When Continue, enter the next iteration
    //When Revert, return to the previous iteration
    //When Exit, save the current iteration and process to skeletonization if (secondChoice == “Continue”) {
        close(“Temp”);
    } else if (secondChoice == “Revert”) {
        close(“Temp”);
close(“Mask_” + stack_count); stack_count--;
     } else {
        close(“Temp”); break;
    }
} else {
break;
   }
}
for (i = 0; i < stack_count; i++) { close(“Mask_” + i);
}
selectWindow(“Mask_” + stack_count);
//Dilation Dialog ends

//Create skeleton, overlay with Binary using imageCalculator rename(“Skeleton”);
run(“Skeletonize”);
imageCalculator(“add create”, “Binary”, “Skeleton”);
close(“Skeleton”);
close(“Binary”);
selectWindow(“Result of Binary”);
rename(“newBinary”);
saveAs(“tiff”, binaryPath);
rename(“newBinary”);

//Soma Analysis
selectWindow(“newBinary”);
run(“Duplicate…”, “ “);
rename(“newBinary2”);
run(“Watershed”);
run(“Clear Results”);
run(“Analyze Particles…”);
n = roiManager(“count”);
somaArray = newArray(n);
for (var i = 0; i < n; i++) {
somaArray[i] = i;
}
selectWindow(“Results”);
saveAs(“Measurements”, somaResultsPath);
run(“Clear Results”);
selectWindow(“newBinary2”);
open(imageOriginPath+”.tif”);
run(“Enhance Contrast”, “saturated=0.35”);
run(“Apply LUT”);
roiManager(“Select”, somaArray);
selectWindow(basename+”.tif”);
setOption(“Show All”, true);
run(“Flatten”);
saveAs(“tiff”,somaOutlinesPath);
run(“Close”);
close(“newBinary2”);
roiManager(“reset”);
//Cell Analysis selectWindow(“newBinary”);
run(“Analyze Particles…”);
n1 = roiManager(“count”);
// Can delete ROIs here
waitForUser(“ROI Deletion”, “In the ROI manager list, delete ROIs (using [Delete]). Click ’OK’ to continue.”);
n2 = roiManager(“count”);
if (n2 < n1) {
run(“Clear Results”);
roiManager(“multi-measure append”);
}
cellArray = newArray(n2);
for (var i = 0; i < n2; i++) { cellArray[i] = i;
}
open(imageOriginPath+”.tif”);
run(“Enhance Contrast”, “saturated=0.35”);
run(“Apply LUT”);
roiManager(“Select”, cellArray);
selectWindow(basename+”.tif”);
setOption(“Show All”, true);
run(“Flatten”);
saveAs(“tiff”,wholeCellOutlinesPath);
roiManager(“XOR”);
run(“Create Mask”);
selectWindow(“Results”);
saveAs(“Measurements”, wholeCellResultsPath);
close(“newBinary”);
run(“Clear Results”);
close(“newBinary”);

//Skeletonize and Analyze Skeletons
selectWindow(“Mask”);
rename(“Skeleton”);
run(“Skeletonize”);
saveAs(“tiff”,skeletonPath);
rename(“Skeleton”);
run(“Analyze Skeleton (2D/3D)”,”prune=[lowest intensity branch] show display”);
selectWindow(“Results”);
saveAs(“Measurements”, skeletonResultsPath);
selectWindow(“Branch information”);
saveAs(“Measurements”, branchInformationPath);
//Close all windows, end macro. close(“*”);
selectWindow(“ROI Manager”);
run(“Close”);
selectWindow(“Results”);
run(“Close”);
if (isOpen(“BranchInformation.csv”)) { selectWindow(“BranchInformation.csv”);
run(“Close”);
}
waitForUser(“Finish”, “Results in the output folder. Click ’OK’ to exit.”);
~~~

## SUPPLEMENTARY NOTE 5

### AUTOMORFI MACRO FOR COLOR PHOTOGRAPHY (IMAGEJ/FIJI-COMPATIBLE)

~~~
//AutoMorFi_3 macro adapted for color photography

close(“*”);
roiManager(“reset”);
run(“Clear Results”); setOption(“BlackBackground”, false);
path = File.openDialog(“Choose a File”);
basename = File.nameWithoutExtension();
dirname = File.getParent(path);
folder = dirname + File.separator + basename; if (File.exists(folder)) {
print(“Directory already exists, override the existing directory”);
list = getFileList(folder);
for (i = 0; i < list.length; i++) {
File.delete(folder + File.separator + list[i]);
// Delete each file
      }
}
else {
File.makeDirectory(folder);
}

binary = folder + File.separator + “Binary”;
objectOutline = folder + File.separator + “ObjectOutline”; objectMeasure = folder + File.separator + “ObjectMeasurement.csv”; somaResultsPath = folder + File.separator + “SomaResults.csv”; somaOutlinesPath = folder + File.separator + “SomaOutlines”; skeletonPath = folder + File.separator + “Skeleton”; skeletonResultsPath = folder + File.separator + “SkeletonResults.csv”;
branchInformationPath = folder + File.separator + “BranchInformation.csv”;

run(“Bio-Formats Importer”, “open=”+path);
imageOriginPath = folder + File.separator + basename; saveAs(“tiff”, imageOriginPath);

getPixelSize(unit, pixelWidth, pixelHeight);
scale = 1 / pixelWidth;
nameOrigin = getTitle();
microWidth = getWidth() / scale;
microHeight = getHeight() / scale;

selectWindow(basename+”.tif”);
run(“Duplicate…”, “ “);
rename(“Binary”);
run(“8-bit”);
run(“Enhance Contrast”, “saturated=0.35”);
run(“Smooth”);
saveAs(“tiff”,binary);
run(“Threshold…”);
waitForUser; close(“Threshold”);
run(“Despeckle”);
run(“Invert LUT”);

stack_count = 0; //Keep track of the number of dilations rename(“Mask_” + stack_count);
while (true) {
//Creating dialog with dilate option “Yes”/”No” Dialog.create(“Erode Option”);
Dialog.addChoice(“Do you want to erode the image?”, newArray(“Yes”, “No”));
Dialog.show();
choice = Dialog.getChoice();
//When choosing “Yes”, dilate the image from the previous stack if (choice == “Yes”) {
selectWindow(“Mask_” + stack_count);
run(“Duplicate…”, “title=Temp”);
run(“Erode”);
stack_count++;
selectWindow(“Temp”);
rename(“Mask_” + stack_count);
//Creating dialog with options “Continue”, “Revert”, “Exit” Dialog.create(“Continue or revert?”);
Dialog.addChoice(“Continue erosion?”, newArray(“Continue”, “Revert”, “Finished”));
Dialog.show();
secondChoice = Dialog.getChoice();
//When Continue, enter the next iteration
//When Revert, return to the previous iteration
//When Exit, save the current iteration and process to skeletonization if (secondChoice == “Continue”) {
close(“Temp”);
} else if (secondChoice == “Revert”) { close(“Temp”);
close(“Mask_” + stack_count);
stack_count--;
} else {
close(“Temp”);
break;
     }
} else {
break;
    }
}
for (i = 0; i < stack_count; i++) { close(“Mask_” + i);
}
selectWindow(“Mask_” + stack_count);
rename(“Binary”);
run(“Duplicate…”, “ “);
rename(“Soma”);
run(“Duplicate…”, “ “);
rename(“Skeleton”);
selectWindow(“Binary”);
run(“Analyze Particles…”);
n1 = roiManager(“count”);
// Can delete ROIs here
waitForUser(“ROI Deletion”, “In the ROI manager list, delete ROIs (using [Delete]). Click ’OK’ to continue.”);
n2 = roiManager(“count”);
if (n2 < n1) {
run(“Clear Results”);
roiManager(“multi-measure append”);
}
objectArray = newArray(n2);
for (var i = 0; i < n2; i++) { objectArray[i] = i;
}

selectWindow(“Results”);
saveAs(“Measurements”, objectMeasure);
close(“Results”);
roiManager(“Select”, objectArray);
selectWindow(basename+”.tif”);
setOption(“Show All”, true);
run(“Flatten”);
saveAs(“tiff”,objectOutline);
run(“Close”);
close(“Binary”);
roiManager(“reset”);
run(“Clear Results”);
selectWindow(“Soma”);
run(“Watershed”);
run(“Analyze Particles…”);
n = roiManager(“count”);
somaArray = newArray(n);
for (var i = 0; i < n; i++) {
somaArray[i] = i;
}
selectWindow(“Results”);
saveAs(“Measurements”, somaResultsPath);
run(“Clear Results”);
selectWindow(“Soma”);
open(imageOriginPath+”.tif”);
run(“Enhance Contrast”, “saturated=0.35”);
roiManager(“Select”, somaArray);

selectWindow(basename+”.tif”);
setOption(“Show All”, true);
run(“Flatten”);
saveAs(“tiff”,somaOutlinesPath);
run(“Close”);
close(“Soma”);
run(“Clear Results”);

selectWindow(“Skeleton”);
run(“Skeletonize”);
saveAs(“tiff”,skeletonPath);
rename(“Skeleton”);
run(“Analyze Skeleton (2D/3D)”,”prune=[lowest intensity branch] show display”);
selectWindow(“Results”);
saveAs(“Measurements”, skeletonResultsPath);
selectWindow(“Branch information”);
saveAs(“Measurements”, branchInformationPath);
close(“*”);
selectWindow(“ROI Manager”);
run(“Close”);
selectWindow(“Results”);
run(“Close”);
~~~

## SUPPLEMENTARY NOTE 6

### CELL MATCHING APP (“ANALYZER”)

AutoMorFi generates “soma” and “whole cell” measurements in separate .csv output files. The cell matching algorithm combines the results by correlating every soma to its whole cell by using their common xy-coordinate system to find the soma with the shortest Euclidean distance to its cell. The algorithm runs on Python using the K-Nearest-Neighbours (KNN) model, which finds the k value of nearest neighbors to a given data point using the Euclidean distance between two points. Notably, in rare instances, a detected soma may be deemed have the shortest distance to a target cell to which the soma does not belong. To ensure an accurate match, an arbitrary default threshold is used to limit matched soma-cell pairs to within 12 µm in horizontal, vertical, and Euclidean distances.

#### Tutorial for Cell Matching App

1. Open Command Prompt
2. Connect to the folder containing the scripts (Supplementary Software 1) cd path/to/your/folder
3. Create the python virtual environment by python -m venv “name” Make sure Python is installed. Check Python version by typing python -V
4. Activate virtual environment by .\name\Scripts\activate

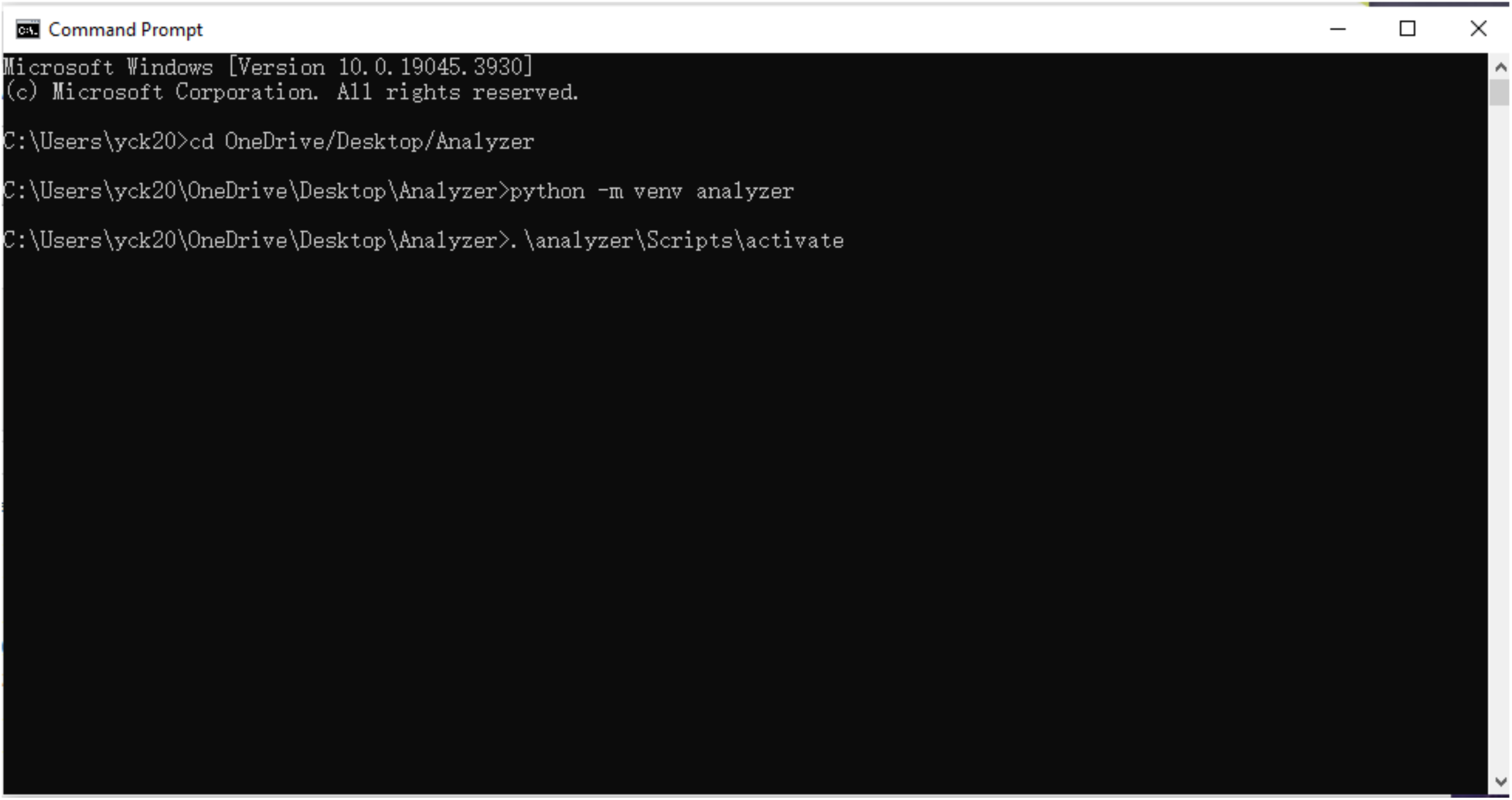

5. When the virtual environment is activated, your command prompt switches to a new window.
6. Use pip or pip3 to install all required libraries including numpy, scipy, scikit-learn, pandas, openpyxl, tk, and tkinterdnd2-universal. Check for all required libraries in **install.sh**. The installation statement is in the first line.

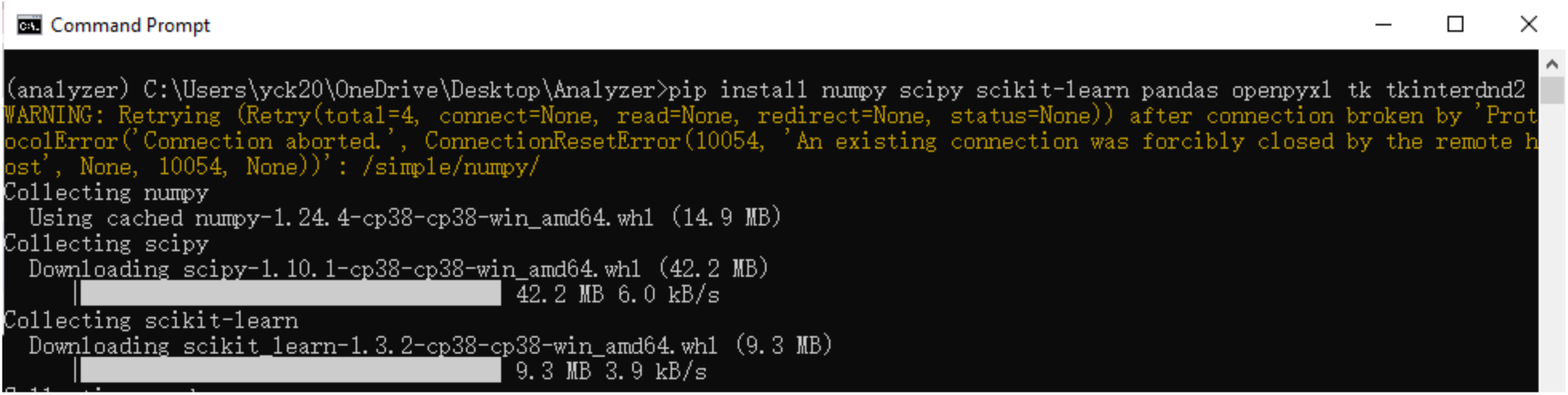

7. When all the libraries are installed, run **python main.py**

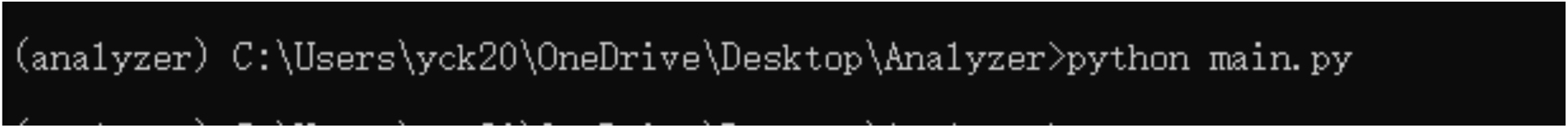

8. For future use, double click **main.py** to open the cell matching app without using the command prompt.

#### Troubleshooting for Cell Matching App

1. Use Python versions 3.7, 3.8, or 3.9.
2. Try different installing commands: pip, pip3, or pip3.x according to your version of Python.
3. Install tkinterdnd2-universal instead of tkinterdnd2 because the latter is incompatible with GitHub branches.
4. Run the code inside a virtual environment.
5. Run the code as the PC administrator.

