## SupplementarySoftware1 for "AutoMorFi: Automated Whole-image Morphometry in Fiji/ImageJ for Diverse Analyses and Discoveries": main.py

```
import os
import tkinter as tk
from tkinter import ttk
from tkinterdnd2 import DND_FILES, TkinterDnD
from match import Analyzer
import pandas as pd

target_files = [
        "SomaResults.csv",
        "WholeCellResults.csv",
        "SkeletonResults.csv",
]


def detect_files(folder_path):

    files = []
    files_dict = {}
    for root, _, filenames in os.walk(folder_path):
        for filename in filenames:
            if filename in target_files:
                file_path = os.path.join(root, filename)
                files.append(file_path)
                name, _ = os.path.splitext(filename)
                files_dict[name] = file_path
    return files, files_dict


def process_folder(folder_path, progress, total_folders):
    print(folder_path)
    file_paths, dict= detect_files(folder_path)

    found_files = [os.path.basename(path) for path in file_paths]
    missing_files = [f for f in target_files if f not in found_files]
    error_message = ""
    unprocessed = ""
    if len(file_paths) == 3:
        soma = dict.get("SomaResults")
        cell = dict.get("WholeCellResults")
        skeleton = dict.get("SkeletonResults")


        analyzer = Analyzer()
        analyzer.addFiles(SomaResults=soma, WholeCellResults=cell, SkeletonResults=skeleton)
        analyzer.Analyze()
        output1 = analyzer.getOutput()
        output2 = analyzer.getCleanOutput()
        y, error = analyzer.getError()
        
        if y is False:
            filename1 = os.path.basename(folder_path) + "_ALL_DATA.csv"
            filename2 = os.path.basename(folder_path) + "_PLOTTING_DATA.csv"
            output1_location = os.path.join(folder_path, filename1)
            output1.to_csv(output1_location, index=False)
            output2_location = os.path.join(folder_path, filename2)
            output2.to_csv(output2_location, index=False)
        else:
            error_message = error
            unprocessed = os.path.basename(folder_path)
            unprocessed_folder.append(unprocessed)


    else:
        print("The following required files are missing:")
        for missing_file in missing_files:
            print("-", missing_file)
        
    current_directory = os.path.dirname(os.path.abspath(__file__))

    update_text_info(current_directory, folder_path, file_paths, error_message,unprocessed_folder)

    progress["value"] += 100/total_folders  
    progress.update()


def on_drop(event):
    received_data = event.data.strip()
    print("Initial received_data:", received_data)
    
    try:
        # List all contents of the received folder, excluding hidden/system files
        folder_contents = [name for name in os.listdir(received_data) if not name.startswith('.')]
        print("Contents of received_data (excluding hidden/system files):", folder_contents)

        # Check if the received folder only contains other folders (subdirectories)
        if all(os.path.isdir(os.path.join(received_data, name)) for name in folder_contents):
            # If so, set received_data to a space-separated list of these folders
            subfolders = [os.path.join(received_data, name) for name in folder_contents]
            print("All subfolders:", subfolders)
            received_data = " ".join(subfolders)
            print("Modified received_data:", received_data)
    except:
        pass

    folder_paths = received_data.split(' ')
    print("Received folder paths:", folder_paths)
    total_folders = len(folder_paths)  # total number of folders
    progress['value'] = 0  # reset progress bar
    unprocessed_folder.clear() #reset
    for folder_path in folder_paths:
 
        process_folder(folder_path.strip(), progress, total_folders)


def update_text_info(directory, folder_path, file_paths, error_message, unprocessed):

    info_text.delete(1.0, tk.END)
    info_text.insert(tk.END, f"Working Directory: {directory}\n")
    info_text.insert(tk.END, f"Folder Path: {folder_path}\n")
    for path in file_paths:
        info_text.insert(tk.END, f"File: {path}\n")
    if error_message != "":
        info_text.insert(tk.END, f"ERROR: {error_message}\n")
    for item in unprocessed:
        info_text.insert(tk.END, f"ERROR FILE: {item}\n")

    


def main():
    root = TkinterDnD.Tk()
    root.title("Autogliji Data Processer")

    root.geometry('800x600')

    global unprocessed_folder
    unprocessed_folder = []

    label = tk.Label(root, text="Drag and drop a folder here:")
    label.pack(pady=10)

    drop_target = tk.Label(root, text="Drop folder here", relief=tk.RAISED, width=80, height=10)
    drop_target.pack(padx=10, pady=10)
    drop_target.drop_target_register(DND_FILES)
    drop_target.dnd_bind('<<Drop>>', on_drop)

    global progress

    style = ttk.Style()
    
    style.configure("TProgressbar", troughcolor='gray', background='blue', thickness=50)

    progress = ttk.Progressbar(root, orient=tk.HORIZONTAL, length=600, mode='determinate', style="TProgressbar")
    progress.pack(pady=20)

    info_frame = tk.Frame(root)
    info_frame.pack(side=tk.BOTTOM, fill=tk.BOTH, padx=10, pady=10)

    global info_text
    info_text = tk.Text(info_frame, width=80, height=20)
    info_text.pack(fill=tk.BOTH, padx=10, pady=10)


    root.mainloop()

if __name__ == "__main__":
    main()
```
