## SupplementarySoftware1 for "AutoMorFi: Automated Whole-image Morphometry in Fiji/ImageJ for Diverse Analyses and Discoveries": match.py

```
import pandas as pd
import numpy as np
from sklearn.neighbors import NearestNeighbors
import os
import warnings

warnings.simplefilter("ignore")

##########  DATA IMPORT  ###########
class Analyzer():
    def __init__(self, thresholdx=12, thresholdy=12, thresholddist=12):
        self.SomaResults = ""
        self.WholeCellResults = ""
        self.SkeletonResults = ""
        self.soma = pd.DataFrame()
        self.cell = pd.DataFrame()
        self.skeleton = pd.DataFrame()
        self.thresholdx = thresholdx
        self.thresholdy = thresholdy
        self.thresholddist = thresholddist
        self.output = pd.DataFrame()
        self.cleanOutput = pd.DataFrame()
        self.warn = False
        self.error = ""


    def addFiles(self, SomaResults, WholeCellResults, SkeletonResults):
        self.SomaResults = SomaResults
        self.WholeCellResults = WholeCellResults
        self.SkeletonResults = SkeletonResults


    def Analyze(self):
        
        def drop_params(df, prefix):
            all_params_to_drop = ["X", "Y", "Slice", "FeretX", "FeretY", "Angle", "IntDen", "RawIntDen", "Median", "XM", "YM", "%Area", "Mean", "StdDev", "Mode", "Min", "Max", "Skew", "Kurt", "BX", "BY", "FeretAngle"]
            all_params_to_drop = [prefix + '_' + param for param in all_params_to_drop]

            params_to_drop = [col for col in df.columns if col in all_params_to_drop]
            df_dropped = df.drop(columns=params_to_drop)
            return df_dropped


        def rename_params(df, prefix):
            df.rename(columns={"...1": "ID"}, inplace=True)
            df.columns = ['ID' if col == df.columns[0] else col for col in df.columns]
            df.columns = [prefix + '_' + col for col in df.columns[0:]]
            return df

        def rename_skeleton_params(df):
            df.rename(columns={"# Branches": "Segment",
                            "# Junctions": "Junction",
                            "# End-point voxels": "End-pt Voxels",
                            "# Slab voxels": "Slab Voxels",
                            "# Triple points": "Triple-pts",
                            "# Quadruple points": "Quad-pts",
                            "Maximum Branch Length": "Maximum Branch Length"}, inplace=True)
            return df
        
        try:
            self.soma = pd.read_csv(self.SomaResults)
            #self.soma = drop_params(self.soma)
            self.soma = rename_params(self.soma, "SOMA")
            self.soma["SOMA_ID"] = self.soma["SOMA_ID"].astype(str)

            self.wholecell = pd.read_csv(self.WholeCellResults)
            #self.wholecell = drop_params(self.wholecell)
            self.wholecell = rename_params(self.wholecell, "WHOLECELL")
            self.wholecell["WHOLECELL_ID"] = self.wholecell["WHOLECELL_ID"].astype(str)

            self.skeleton = pd.read_csv(self.SkeletonResults)
            self.skeleton.columns = ['WHOLECELL_ID'] + list(self.skeleton.columns[1:])
            self.skeleton = rename_skeleton_params(self.skeleton)
            self.skeleton["WHOLECELL_ID"] = self.skeleton["WHOLECELL_ID"].astype(str)


            ##############  EXTRACT [X, Y, ID]  ##############
            soma_df = self.soma[["SOMA_X", "SOMA_Y", "SOMA_ID"]]
            wholecell_df = self.wholecell[["WHOLECELL_X", "WHOLECELL_Y", "WHOLECELL_ID"]]

            ###  MATCH WHOLECELL x SOMA  ###              
            ##############  CREATE KNN WHERE N=1  ##############
            nbrs = NearestNeighbors(n_neighbors=1).fit(soma_df[["SOMA_X", "SOMA_Y"]].values)
            distances, indices = nbrs.kneighbors(wholecell_df[["WHOLECELL_X", "WHOLECELL_Y"]].values)

            ##############  COMBINE MATCHED COORDINATES  ##############
            wholecell_soma = wholecell_df
            wholecell_soma["temp"] = indices[:, 0]
            wholecell_soma = wholecell_soma.merge(soma_df, left_on='temp', right_index=True, how='left')
            wholecell_soma = wholecell_soma.drop(columns=["temp"])

            ##############  FILTER BY THRESHOLD  ##############
            dx = wholecell_soma["WHOLECELL_X"] - wholecell_soma["SOMA_X"]
            dy = wholecell_soma["WHOLECELL_Y"] - wholecell_soma["SOMA_Y"]
            distances = np.sqrt(dx**2 + dy**2)
            in_threshold_wholecell_soma = wholecell_soma[(abs(dx) <= self.thresholdx) & (abs(dy) <= self.thresholdy) & (distances <= self.thresholddist)]
            unmatched_wholecell_A = wholecell_df[~wholecell_df["WHOLECELL_ID"].isin(in_threshold_wholecell_soma["WHOLECELL_ID"])]
            unmatched_soma = soma_df[~soma_df["SOMA_ID"].isin(in_threshold_wholecell_soma["SOMA_ID"])]

            ##############  FILLING EMPTY AREA FOR UNMATCHED  ##############
            unmatched_wholecell_A["SOMA_ID"] = 0
            unmatched_wholecell_A["SOMA_X"] = 0
            unmatched_wholecell_A["SOMA_Y"] = 0
            unmatched_soma["WHOLECELL_ID"] = 0
            unmatched_soma["WHOLECELL_X"] = 0
            unmatched_soma["WHOLECELL_Y"] = 0

            self.output = pd.concat([in_threshold_wholecell_soma, unmatched_wholecell_A, unmatched_soma], ignore_index=True)
            self.output = self.output.drop(columns=["temp", "SOMA_X", "SOMA_Y", "WHOLECELL_X", "WHOLECELL_Y"])
            self.output = self.output.merge(self.soma, on=['SOMA_ID'], how='outer')
            self.output = self.output.merge(self.wholecell, on=['WHOLECELL_ID'], how='outer')
            self.output = self.output.merge(self.skeleton, on=['WHOLECELL_ID'], how='outer')
            #self.output = self.output.drop(columns=["SOMA_X", "SOMA_Y", "WHOLECELL_X", "WHOLECELL_Y"])
            self.output.fillna(0, inplace=True)
            self.output["Total Branch Length"] = self.output["Segment"] * self.output["Average Branch Length"]
            self.output["Average Branch Length"] = self.output["Total Branch Length"] / self.output["End-pt Voxels"]
            self.output["Cell Spread"]  = self.output["WHOLECELL_AR"] / self.output["End-pt Voxels"]
            self.output["ImageID"] = os.path.basename(os.path.dirname(self.SomaResults))
            self.output['condition'] = None
            temp = ['ImageID', 'condition']
            cols = temp + [col for col in self.output if col not in temp]
            self.output = self.output[cols]

            if self.output is None:
                self.warn = True
                e = "All_Data DataFrame is None"
            


            self.cleanOutput = self.output[~((self.output['WHOLECELL_ID'] == 0) | (self.output['SOMA_ID'] == 0))]

            # Add a safety check to drop rows if ROI does not contain skeletonization analysis data:
            self.cleanOutput = self.cleanOutput[~(self.cleanOutput['End-pt Voxels'] == 0)]

            self.cleanOutput = drop_params(self.cleanOutput, "WHOLECELL")
            self.cleanOutput = drop_params(self.cleanOutput, "SOMA")


        except Exception as e:
            self.warn = True
            self.error = e
    

    def getOutput(self):
        return self.output
    
    def getCleanOutput(self):
        return self.cleanOutput
    
    def getError(self):
        if self.warn is False :
            return False, self.error
        else:
            return True, self.error
```
